# DNA-directed termination of mammalian RNA polymerase II

**DOI:** 10.1101/2024.06.01.596947

**Authors:** Lee Davidson, Jérôme O Rouvière, Rui Sousa-Luís, Takayuki Nojima, Nicholas Proudfoot, Torben Heick Jensen, Steven West

## Abstract

The best-studied mechanism of eukaryotic RNA polymerase II (RNAPII) transcriptional termination involves polyadenylation site-directed cleavage of the nascent RNA. The RNAPII-associated cleavage product is then degraded by XRN2, dislodging RNAPII from the DNA template. In contrast, prokaryotic RNAP and eukaryotic RNAPIII often terminate directly at T-tracts in the coding DNA strand. Here, we demonstrate a similar and omnipresent capability for mammalian RNAPII. XRN2- and T-tract-dependent termination are independent - the latter usually acting when XRN2 cannot be engaged. We show that T-tracts terminate snRNA transcription, previously thought to require the Integrator complex. Importantly, we find genome-wide termination at T-tracts in promoter-proximal regions, but not within protein-coding gene bodies. XRN2-dependent termination dominates downstream of protein-coding genes, but the T-tract process is sometimes employed. Overall, we demonstrate global DNA-directed attrition of RNAPII transcription, suggesting that RNAPs retain the potential to terminate over T-rich sequences throughout evolution.

## INTRODUCTION

Transcriptional termination of mammalian RNAPII has historically been studied at long protein-coding genes. Here, the process is coupled to 3’ end cleavage and polyadenylation (CPA) of the nascent RNA, which occurs immediately downstream of a polyadenylation site (PAS) (Eaton and West 2020; Rodriguez-Molina et al. 2023). The RNAPII-associated 3’ cleavage product is then degraded 5’→3’ by the XRN2 exonuclease, resulting in transcriptional termination (Eaton et al. 2020; Eaton and West 2020; Rodriguez-Molina et al. 2023). This is further facilitated by dephosphorylation of the elongation co-factor, SPT5, by the PNUTS-PP1 phosphatase (Cortazar et al. 2019; Eaton et al. 2020). Similar “torpedo” mechanisms are employed in lower eukaryotes, prokaryotes, and archaea (Sikova et al. 2020; Wiedermannova and Krasny 2021). Some of these involve endonucleolytic cleavage and exoribonucleolytic activities (Sanders et al. 2020). In other cases, the Rho RNA helicase tracks along the RNAP-associated transcript to elicit termination (Molodtsov et al. 2023).

Other functional mammalian transcripts also undergo 3’ end processing by endonucleolytic cleavage. Small nuclear (sn)RNAs are critical components of the spliceosome, which removes introns from pre-messenger (m)RNA (Sanford and Caceres 2004). They possess a so-called 3’ box RNA element, which is recognised by the Integrator (INT) complex and cleaved by its resident endonuclease, INTS11, serving as an snRNA maturation factor (Baillat et al. 2005; Guiro and Murphy 2017). INT also associates with phosphatase activity, through the interaction between its INTS6 subunit and the PP2A complex (Zheng et al. 2020). In addition to its role in snRNA processing, INT attenuates transcription in the vicinity of most human RNAPII promoters and an important function of this activity is to suppress antisense and otherwise cryptic transcription (Elrod et al. 2019; Tatomer et al. 2019; Lykke-Andersen et al. 2021; Stein et al. 2022). INT is suggested to act early in transcription by targeting RNAPII that is paused in proximity to the transcription start site (TSS) (Beckedorff et al. 2020; Stein et al. 2022). Structural and molecular studies support this by showing that INT forms a complex with RNAPII associated with negative elongation factor (NELF) – a mediator of promoter-proximal pausing (Stadelmayer et al. 2014; Yamamoto et al. 2014; Fianu et al. 2021). In the structure, the endonuclease active site of INTS11 is open and available, allowing cleavage when the nascent RNA is exposed. Depletion of INT components increases transcription beyond snRNA loci, which has been interpreted to reflect a termination mechanism analogous to that at protein-coding genes (Baillat et al. 2005; Stadelmayer et al. 2014; Yamamoto et al. 2014). The model predicts that INTS11-dependent cleavage at the 3’ box exposes an RNAPII-associated RNA to 5’→3’ degradation by XRN2. However, XRN2 has no apparent impact on snRNA transcriptional termination, suggesting either the involvement of a different 5’→3’ exonuclease or an alternative mechanism (Eaton et al. 2018).

Replication-dependent histone (RDH) genes also produce short and intronless RNA. These RNAPII-dependent RNAs are transcribed from RDH gene clusters during S-phase (Marzluff and Koreski 2017). Although the CPSF73 endonuclease processes their 3’ ends, RDH RNAs are not polyadenylated, unlike other mRNAs (Dominski et al. 2005; Sun et al. 2020). Further contrasting with conventional mRNA 3’ end formation, the RDH processing machinery is recruited by the U7 snRNA rather than *via* a PAS (Yang et al. 2020). RDH 3’ end formation occurs co-transcriptionally and there is some evidence that XRN2 can target the RNAPII-associated cleavage product (Sousa-Luis et al. 2021; Cortazar et al. 2022). However, XRN2 has a more limited impact on the termination of RDH transcription compared to that of canonical protein-coding genes (Fong et al. 2015; Eaton et al. 2018). Therefore, additional uncharacterised mechanisms may be used to ensure efficient transcriptional termination.

Beyond the gene classes outlined above, mammalian genomes are pervasively transcribed, yielding a variety of short and largely non-coding (nc) transcripts (Chen et al. 2016). Promoter upstream transcripts (PROMPTs) are a prominent example of this and are transcribed upstream and antisense of most protein-coding TSSs (Preker et al. 2008; Preker et al. 2011). PROMPT transcription is generally terminated within approximately 3 kilobases (kb) of their TSSs and the resulting RNAs are rapidly degraded in a 3’→5’ direction by the nuclear exosome (Preker et al. 2008; Davidson et al. 2019). Similar short and rapidly degraded transcripts are produced in the sense direction of protein-coding genes, which reveals widespread early attrition of RNAPII transcription (Iasillo et al. 2017; Ogami et al. 2017; Davidson et al. 2019). The mechanisms of such termination are beginning to be uncovered and utilise multiple processes that target immature transcription elongation complexes (Nojima and Proudfoot 2022; Rodriguez-Molina et al. 2023; Wagner et al. 2023). These include INT and the recently described Restrictor complex, which limit the extent of PROMPT and other cryptic transcription (Austenaa et al. 2021; Estell et al. 2021; Estell et al. 2023; Rouviere et al. 2023). Furthermore, PASs can mediate promoter-proximal transcription termination when the elongation-promoting U1 snRNA is not recruited to pre-mRNA (Kaida et al. 2010; Oh et al. 2017).

A distinguishing feature of most mammalian protein-coding genes *vs.* most other transcription units (TUs) is their length, which frequently exceeds 100kb due to long intronic sequences. Their full-length transcription therefore requires robust RNAPII elongation complexes, capable of lengthy transcription of nucleosome-containing templates while simultaneously supporting co-transcriptional RNA processing. The maturation of RNAPII to achieve such elongation competence appears to be a gradual process that occurs over an initial “pausing zone” of ∼3kb (Fong et al. 2022). Within this zone, RNAPII is prone to pausing and premature transcriptional termination by mechanisms including those mentioned above (Lykke-Andersen et al. 2021). Indeed, exosome depletion stabilises short (<3kb) promoter-proximal transcripts at thousands of protein-coding genes (Davidson et al. 2019; Lykke-Andersen et al. 2021). Cyclin-dependent kinase 9 (CDK9) activity promotes RNAPII escape from the pausing zone, presumably through its role in phosphorylating the RNAPII C-terminal domain (CTD) and SPT5 (Jonkers and Lis 2015; Fujinaga et al. 2023). This accelerates RNAPII transcription through the protein-coding gene body (Jonkers et al. 2014). After crossing the PAS, the transcription complex once again reverts to a slower termination-prone state triggered by the dephosphorylation of SPT5 by PNUTS-PP1 (Parua et al. 2018; Cortazar et al. 2019; Eaton et al. 2020). These observations suggest that RNAPII is most vulnerable to termination when situated promoter-proximally or downstream of a PAS.

DNA sequence plays a direct role in transcriptional termination in many biological contexts. Bacterial RNAP often undergoes transcriptional termination by an intrinsic mechanism (Gusarov and Nudler 1999), which involves the transcription of a hairpin structure followed by a U-tract (representing T’s in the coding DNA strand). This results in a thermodynamically weak dA: rU hybrid within RNAP, favouring termination (Martin and Tinoco 1980). The upstream hairpin aids the process possibly by forward translocation of the RNAP, by shearing of the dA: rU hybrid, or by promoting RNAP conformational change. An analogous mechanism applies for eukaryotic RNAPIII, which terminates at runs of ≥4 T’s (Arimbasseri et al. 2013; Arimbasseri et al. 2014). Recently, the termination of budding yeast RNAPII downstream of the PAS was also shown to involve T-tracts (Han et al. 2023). This was further proposed to aid torpedo-dependent termination by the yeast 5’→3’ exonuclease, Rat1. AT-rich regions were also suggested to favour transcriptional termination in mammals (White et al. 2012; Vlaming et al. 2022). Importantly, some T-tracts are sufficient to terminate RNAPII *in vitro*, demonstrating factor-independent activity (Dedrick et al. 1987; Reines et al. 1987). Yet, while termination in most of these examples occurs within a few kb of the TSS, reminiscent of the mammalian pausing zone, any generality and the location of such transcriptional termination elements remains unexplored in higher eukaryotes.

Here, we report the termination of human RNAPII transcription over T-rich sequences. This mechanism is widespread promoter-proximally and over short TUs, including snRNA-, independent snoRNA-, and RDH genes. At snRNA loci, this DNA-directed process is the primary mechanism and transcriptional termination is uncoupled from snRNA 3’ end processing by the INT complex. DNA-directed termination is largely independent of the XRN2-dependent torpedo mechanism and predominantly functions promoter-proximally. However, both processes are used downstream of a subset of protein-coding PASs, where the DNA-directed process can terminate transcription that does not efficiently engage the torpedo mechanism. In contrast, T-tracts are ineffective within the bodies of long protein-coding genes, suggesting that robust RNAPII elongation complexes acquire resistance to this mechanism. Taken together, we identify a vulnerability of immature RNAPII elongation complexes, usually found at the beginning and the end of genes, to terminate at T-rich elements.

## RESULTS

### What is the role of the INT complex in snRNA transcription?

The mechanism of snRNA transcription termination is assumed to resemble that occurring at protein-coding genes because both employ analogous 3’ end processing complexes. Indeed, multiple studies show that INT depletion leads to more transcription downstream of snRNA genes (Baillat et al. 2005; O’Reilly et al. 2014; Davidson et al. 2020; Lykke-Andersen et al. 2021). However, our previous experiments suggested differences because XRN2 did not impact snRNA transcription (Eaton et al. 2018). To better understand what governs the susceptibility of transcription to INT, we reanalysed our previously generated transient transcriptome sequencing (TT-seq) data from control (siCTRL) and INTS11-depleted (siINTS11-treated) HeLa cells (Lykke-Andersen et al. 2021). For every expressed transcript, regardless of its biotype (n=23145), we plotted the TSS-proximal log_2_ fold signal change (siINTS11 *vs.* siCTRL) as a function of the promoter-proximal TT-seq coverage in the siCTRL sample. As previously demonstrated (Lykke-Andersen et al. 2021; Hu et al. 2023), the RNAs with the strongest upregulation following INTS11 depletion tended to derive from lowly expressed TUs (Figure 1A). However, most snRNAs formed clear outliers to this trend by being both highly expressed and sensitive to INTS11 loss.

**FIGURE 1:**
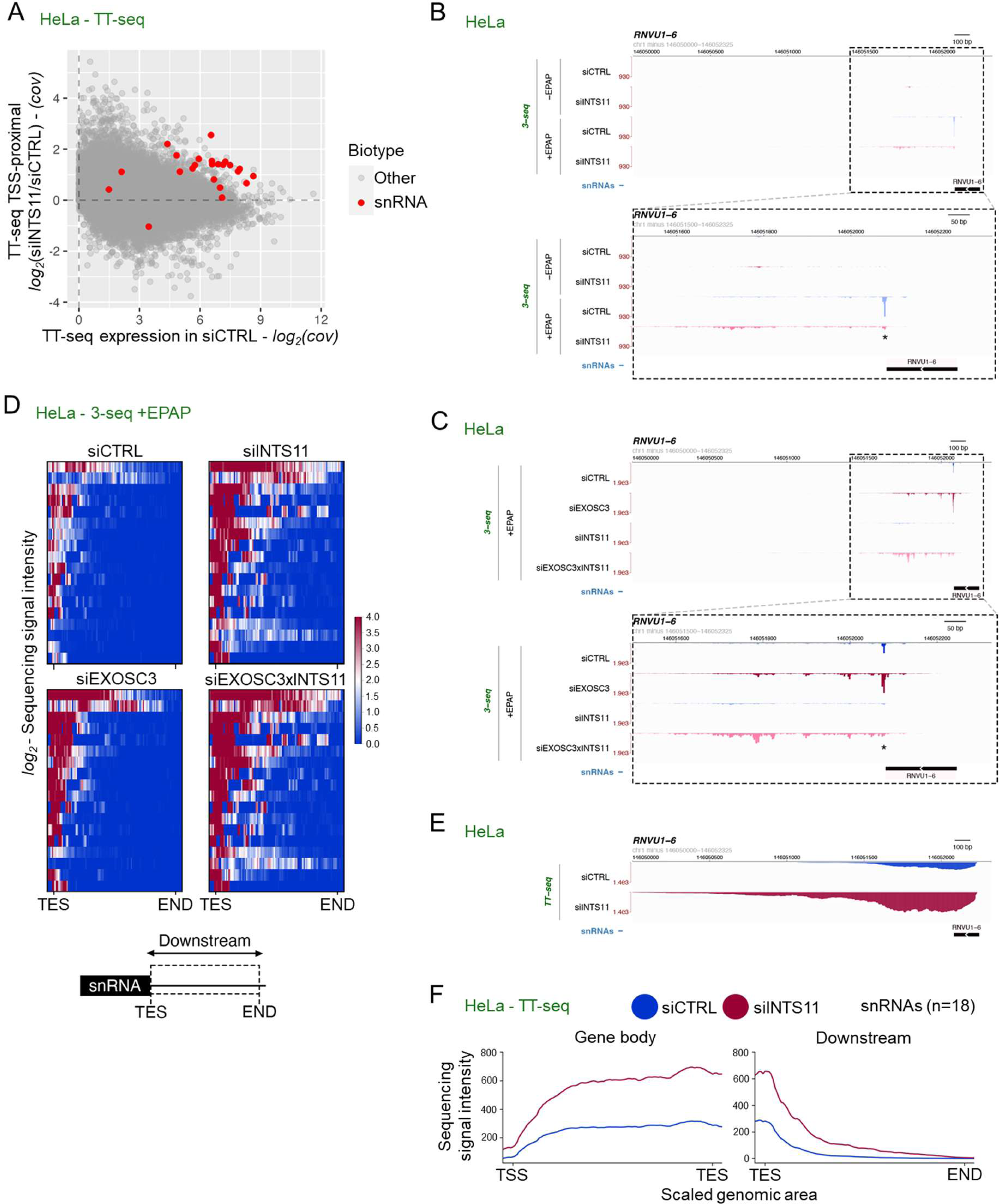
INTS11 is required for snRNA processing but not transcription termination. A. Scatter plot showing the TSS-proximal log_2_ TT-seq signal ratio (siINTS11/siCTRL) (y-axis) as a function of transcript expression levels (x-axis). snRNAs are highlighted in red. The data are from GSE151919 (Lykke-Andersen et al. 2021). B. Genome browser view across and downstream of the *RNVU1-6* TU, displaying RNA 3’ end sequencing (3’-seq) data from siCTRL- and siINTS11-treated HeLa cells (GSE151919 (Lykke-Andersen et al. 2021)). Both -EPAP and +EPAP samples are shown. The top panel encompasses 2kb downstream of the *RNVU1-6* body and the bottom panel shows a zoom-in (dashed box) of the snRNA proximal region with the INTS11 cleavage site indicated by an asterisk. Y-axes display reads per kilobase per million mapped reads (RPKM). C. Genome browser view as in (B) but displaying samples from siCTRL-, siINTS11-, siEXOSC3- or siINTS11xEXOSC3-treated HeLa cells (GSE151919 (Lykke-Andersen et al. 2021)). D. Heatmaps of 3’-seq signal coverage downstream of the transcript end sites (TESs) of 18 snRNAs expressed in HeLa cells. Samples were derived from siCTRL, siINTS11, siEXOSC3 or siINTS11xsiEXOSC3 cells (GSE151919 (Lykke-Andersen et al. 2021)) and RNA was EPAP-treated. The heat scale represents log_2_ sequencing signal intensity from each depletion condition *vs.* the siCTRL. The “END” of this region represents the last position of TT-seq read detection following siEXOSC3 depletion (see Materials and Methods). E. Genome browser view as in (B) but displaying TT-seq data from siCTRL- and siINTS11-treated HeLa cells (GSE151919 (Lykke-Andersen et al. 2021)). F. Metaplot of TT-seq data from siCTRL- or siINTS11-treated HeLa cells (GSE151919 (Lykke-Andersen et al. 2021)) over 18 snRNA loci. The left plot shows snRNA gene body signals from the TSS to the TES, while the right plot shows signals downstream of the TES.

In current models, INT is a productive factor that cleaves and processes pre-snRNAs at their 3’ boxes, which is thought to be coupled to transcriptional termination. This contrasts with the transcriptional attenuation function of INT found elsewhere, which normally leads to RNA decay rather than maturation (Elrod et al. 2019; Tatomer et al. 2019; Lykke-Andersen et al. 2021), and could be why snRNAs were distinguished from most other INT-sensitive transcripts. Alternatively, INT might also attenuate TSS-proximal snRNA transcription upstream of the 3’ box. The consequent increase in transcription that this would cause might appear as increased read-through snRNA transcription following INT loss. These considerations necessitated a re-evaluation of INT function(s) in snRNA biogenesis. Hence, we reanalysed our published sequencing data, representing newly produced RNA 3’ ends (3’-seq) from siCTRL- and siINTS11-treated HeLa cells (Lykke-Andersen et al. 2021). As for the TT-seq data, RNA was metabolically labelled with 4-thiouridine (4sU) and purified by subsequent biotinylation and streptavidin capture (Wu et al. 2020). The isolated nascent RNAs represented a mixture of naturally adenylated and non-adenylated transcripts, with the latter including transcripts isolated from the RNAPII active site and possible transcriptional termination products. In our approach, the 3’ ends of such cellular RNAs lacking a natural poly(A) tail were detected *via* their *in vitro* polyadenylation by *Escherichia coli* poly(A) polymerase (EPAP) (Wu et al. 2020). As 3’ ends of neither mature snRNAs nor their precursors possess a natural poly(A) tail, their detection should be EPAP-dependent.

Illustrating this, the *RNVU1-6* locus displayed almost no signal in samples not subjected to EPAP treatment (-EPAP). However, EPAP treatment (+EPAP) revealed a dominant signal deriving from 3’ box cleavage and lower-level downstream RNA 3’ ends (Figure 1B, note lower panel zoom-in of dashed box). Consistent with INT-dependent snRNA processing, INTS11 depletion abolished the dominant 3’ end. Concomitantly, levels of downstream 3’ ends increased (Figure 1B, compare pink *vs.* blue signals). These 3’ ends could derive from the active site of RNAPII, thereby signifying an anticipated termination defect triggered by INTS11 depletion. Alternatively, they could correspond to post-transcriptional RNAs released from RNAPII terminating at these sites INTS11-independently. To distinguish these possibilities, we analysed RNA 3’-seq data deriving from samples where the RNA exosome subunit EXOSC3 had been individually depleted (siEXOSC3) or co-depleted with INTS11 (siEXOSC3xINTS11). Our rationale was that EXOSC3 depletion would preferentially stabilise 3’ ends of post-transcriptional RNAs whereas nascent 3’ ends would be protected within DNA template-bound RNAPII. Indeed, siEXOSC3 treatment resulted in the accumulation of 3’ ends downstream of the 3’ box (Figure 1C). Moreover, co-depleting INTS11 and EXOSC3 further amplified the signals, suggesting that INTS11-depletion might generally increase snRNA transcription but that the INT complex is not directly required to generate downstream 3’ ends. While co-depleting INTS11 and EXOSC3 increased downstream 3’ end signals, many of their positions were remarkably unchanged between experimental conditions (Figure 1C, lower panel). These salient features observed at the *RNVU1-6* locus were evident for other snRNA TUs and more generally observed by meta-analysis (Figures S1A and Figure 1D). This implies that a significant fraction of snRNA transcriptional termination occurs at specific positions downstream of the snRNA 3’ box *via* an INT-independent process that releases RNA for degradation by the exosome.

Even though the positioning of RNA 3’ ends downstream of snRNA 3’ boxes was largely unaffected by INTS11 depletion, the complex still affects snRNA transcription since these 3’ ends accumulated upon INTS11 depletion. This would be consistent with previous reports on increased transcription downstream of snRNAs when INT is depleted (Baillat et al. 2005; O’Reilly et al. 2014; Davidson et al. 2020; Lykke-Andersen et al. 2021). To reconcile our findings with these observations, we turned to TT-seq samples performed after siCTRL- or siINTS11-treatment of HeLa cells and plotted sequence coverage over and downstream of snRNA loci. In line with the RNA 3’-seq data, the *RNVU1-6* locus displayed an increased TT-seq signal downstream of the 3’ box upon INTS11 depletion (Figure 1E). Although this could be interpreted as a transcriptional termination defect, there was also an increased TT-seq signal over the *RNVU1-6* gene body, suggesting that INTS11 depletion increases snRNA transcription as we suggested above. This was observed for other individual snRNAs (Figure S1B) and by meta-analysis (Figure 1F). A plausible interpretation of these data is that INTS11 depletion increases snRNA transcription without affecting its natural termination mechanism. In this sense, INT also plays an attenuating role at snRNA TUs analogous to its function throughout the genome and its specific role in 3’ box processing.

### INT-driven snRNA 3’ end processing is co-transcriptional

Our analyses challenged the long-standing view that snRNA transcriptional termination requires INT. This model was, at least partly, based on the similarity between snRNA 3’ end processing by INT and PAS-dependent mRNA maturation by the CPA complex. Co-transcriptional cleavage near the PAS is a critical feature of the termination mechanism at protein-coding genes (Eaton and West 2020). Regarding 3’ box processing, our data in Figure 1 demonstrated its occurrence on newly transcribed RNA, but the 4sU labelling approach cannot differentiate between co- and post-transcriptional INT activity. To ask whether INT processes 3’ box-containing RNA co-transcriptionally, we employed POINT5-seq (Sousa-Luis et al. 2021). In this method, chromatin-associated RNAPII is immunopurified with its nascent transcript. The 5’ ends of these RNAs are then mapped with single-nucleotide resolution *via* strand-switching reverse transcription followed by RNA-seq. In parallel, we performed POINT-seq analysis where RNAs from chromatin-associated RNAPII are sequenced in their entirety to provide an overall picture of nascent transcription. These analyses were performed in HCT116 cells in which the INT backbone subunit, INTS1, was tagged with an auxin-inducible degron allowing its depletion within 3hr of auxin (IAA) addition (Figure 2A). This rapid depletion provided confidence that any observed effect was a primary consequence of INTS1 loss.

**FIGURE 2:**
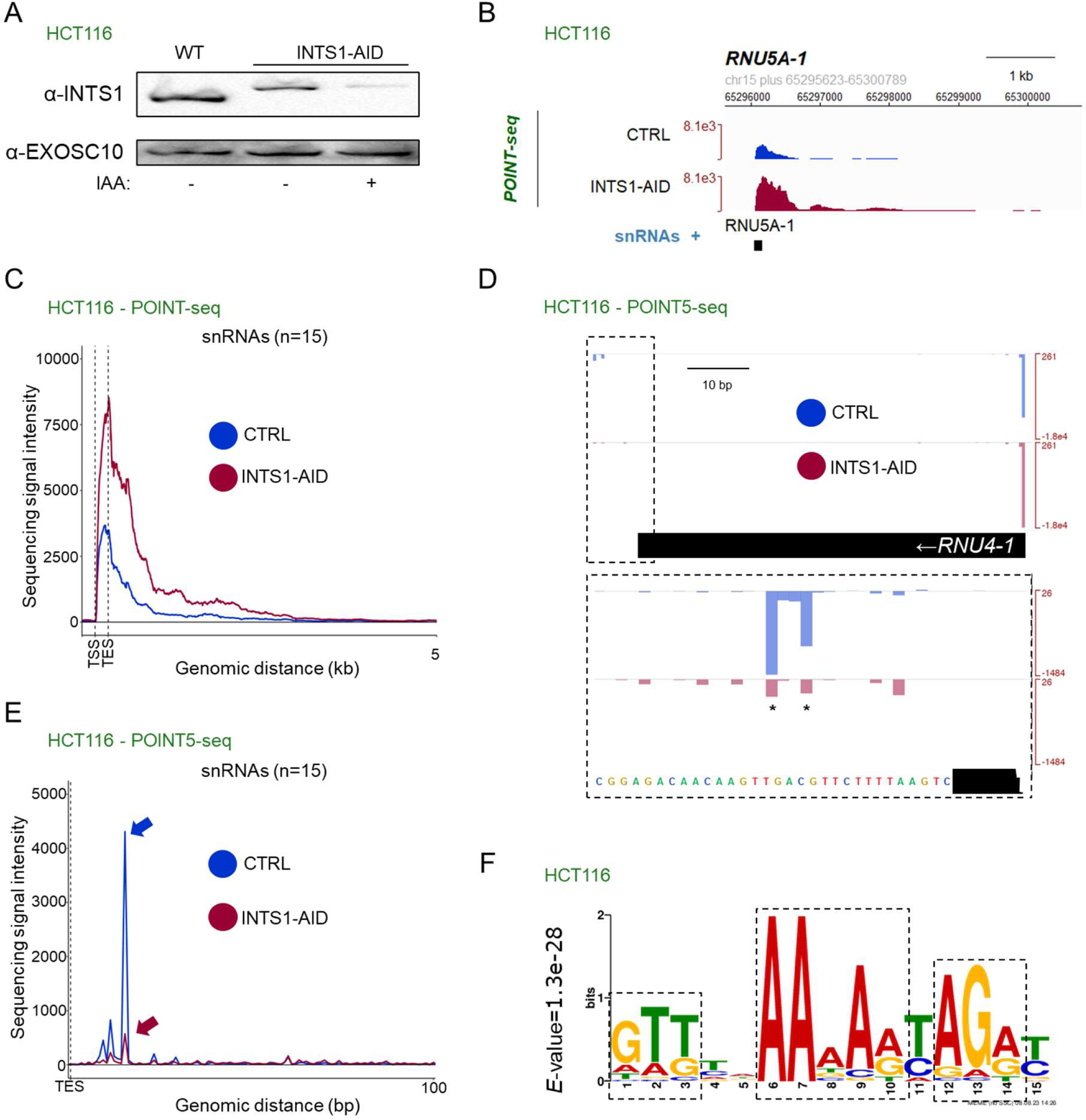
3’ end processing of snRNA precursors occurs co-transcriptionally. A. Western blotting analysis of protein extracts from unmodified (WT) HCT116 cells and *INTS1-AID* HCT116 cells treated (+) or untreated (-) with auxin for 3hrs. Note that degron-tagging increases the INTS1 molecular weight as expected. EXOSC10 was probed as a loading control. B. Genome browser view across and downstream of the *RNU5A-1* TU, displaying POINT-seq data from *INTS1-AID* HCT116 cells untreated (CTRL) or treated (INTS-AID) with auxin. The y-axis sequence signal intensity units are Bins Per Million mapped reads (BPM). C. Metaplot of POINT-seq data of RNA from *INTS1-AID* HCT116 cells untreated (CTRL) or (INTS-AID) with auxin. The plot represents 15 snRNAs separated from their neighbouring TUs by at least 5kb. The x-axis shows from 0.2kb upstream to 5kb downstream of the annotated snRNA (TSS to TES). The y-axis sequence signal intensity units are BPM. D. Genome browser view as in (B), but displaying POINT5-seq data across and downstream of the *RNU4-1* TU. The top panel shows the major TSS peak and the minor downstream INT processing site (dashed box). The lower panel displays increased signal resolution around the TES downstream region. The sites of significantly downregulated POINT5 coverage, upon INTS1 depletion, are marked by asterisks. E. Metaplot as in (C) but displaying POINT5-seq data from snRNA TESs to a region 100bp downstream. The y-axis sequence signal intensity units are BPM. Arrows indicate the major INTS1-sensitive site downstream of the annotated snRNA 3’ end. F. 3’ box consensus element derived from POINT5-seq. INT cleavage occurs immediately upstream of this sequence. The E-value represents the probability of encountering the same sequence by chance. See also Figure S2B.

INTS1-AID depletion caused an elevated POINT-seq signal over the snRNA gene body and throughout the downstream region, exemplified by the *RNU5A-1* TU (Figure 2B). However, the POINT-seq signal eventually returned to background levels even when INTS1 was depleted. Meta-analysis confirmed the generality of this observation for other snRNAs (Figure 2C), while also demonstrating that although the INTS1-AID depletion curve displayed a higher signal than the CTRL curve, they both had a similar shape. This agreed with our conclusion from Figure 1, that INT loss increases snRNA transcription but may not affect the utilisation of major transcriptional termination sites downstream of the TU. The data also demonstrated consistency between siRNA-mediated depletion of INTS11 from HeLa cells and AID degron-mediated depletion of INTS1 from HCT116 cells.

Next, we used the POINT5-seq data to assess whether snRNA 3’ end processing is co-transcriptional. At the exemplary *RNU4-1* locus, POINT5-seq revealed two nascent 5’end signals: a major one at the TSS and another minor doublet just beyond the annotated gene body (Figure 2D, note dashed box highlighting the minor doublet). We assumed the latter represents 3’ box processing by INT because it was reduced upon INTS1 depletion. An additional example of such co-transcriptional processing by INT was provided by the *RNU12* locus (Figure S2A) and was generally evident at other snRNA loci (Figure 2E). These INT cleavage sites were situated downstream of the annotated snRNA 3’ end, consistent with previous findings that mature snRNA 3’ ends are formed by TOE1-mediated 3’→5’ trimming of a remaining short 3’ extension (Lardelli and Lykke-Andersen 2020). Thus, like PAS cleavage, 3’ box processing is co-transcriptional. However, unlike PAS cleavage, eliminating 3’ box processing does not prevent transcriptional termination.

By performing a MEME analysis of the 30 nucleotides immediately downstream of snRNA TESs, we identified a 3’ box consensus element downstream of the detected INT cleavage sites (Figure 2F, S2B). To test its effectiveness as an snRNA 3’ end processing signal, we used a previously established GFP reporter assay (Albrecht and Wagner 2012; Albrecht et al. 2018). In this system, 3’ end box processing precludes GFP expression, which only occurs when processing fails (Figure S2C). By this approach, the consensus 3’ box proved an efficient snRNA processing element (Figure S2D). Interestingly, mutation of its most highly conserved nucleotides had only modest effects on snRNA processing implying that the 3’ box is more resistant to mutation than the PAS, which can be inactivated by single nucleotide substitutions (Proudfoot 2011). We then asked whether similar 3’ box sequences might be active in other promoter-proximal regions. If this was the case, any POINT5-seq signal should be reduced by INTS1 depletion. However, despite being highly active at snRNA loci, the 3’ box consensus remained inactive in other promoter-proximal contexts (Figure S2E). We presume that 3’ box processing requires an snRNA-specific factor or feature, as suggested by prior work showing that snRNA maturation requires transcription from an snRNA promoter (Hernandez and Weiner 1986). Overall, we conclude that 3’ box processing is co-transcriptional and that it is specific for snRNAs.

### The 3’ box elicits snRNA processing but is dispensable for transcription termination

Our above analyses suggested that INT is not critical for snRNA transcription termination. Still, 3’ box cleavage is co-transcriptional and strongly impacts GFP levels in our reporter assay. We therefore more directly addressed whether the 3’ box plays any role in snRNA transcription. Accordingly, we generated two reporter plasmids based on the design from Figure S2C; one containing the U7 snRNA and its 3’ box (pWT) and another from which the 3’ box had been deleted (pΔ3’box) (Figure 3A). Both plasmids contained the downstream GFP gene, the PAS of which we removed to isolate any effect of 3’ box mutation on transcription termination. pWT and pΔ3’box plasmids were transfected into our recently described *INTS11-dTAG* cell line (Eaton et al. 2023) and, following depletion of INTS11, qRT-PCR analysis was performed to assay RNA spanning the 3’ box (UC 3’ box amplicon) or at a downstream read-through position (RT amplicon) (Figure 3A, see positions of utilised amplicons). Deletion of the 3’ box resulted in significantly more RNA spanning the 3’ box region consistent with its processing activity (Figure 3B left panel, compare ‘UC 3’ box’ light blue and pink columns). INTS11 depletion also increased UC 3’ box RNA levels but only for pWT, demonstrating that 3’ box deletion eliminated both 3’ end processing activity and INTS11-sensitivity. Even so, read-through RNA levels were unchanged between pWT and pΔ3’ box constructs in CTRL samples (Figure 3B, compare ‘RT’ light blue and pink columns), suggesting that the 3’ box is unnecessary for transcriptional termination. This contrasts with the widely observed abolition of transcriptional termination on similar protein-coding reporter plasmids containing mutated PASs (Whitelaw and Proudfoot 1986; Dye and Proudfoot 1999; Dye and Proudfoot 2001). Interestingly, INTS11 depletion significantly increased RT RNA levels from both constructs, demonstrating that the INT complex impacts reporter transcription in a 3’ box-independent manner. This reconciles our described INT-independent snRNA transcription termination and the increased snRNA transcription downstream of the TES widely seen following INT loss (including the TT- and POINT-seq data above). Therefore, we suggest that snRNA transcriptional read-through, following INTS11 depletion, derives from increased transcription due to a diminished attenuating activity upstream of the 3’ box.

**FIGURE 3:**
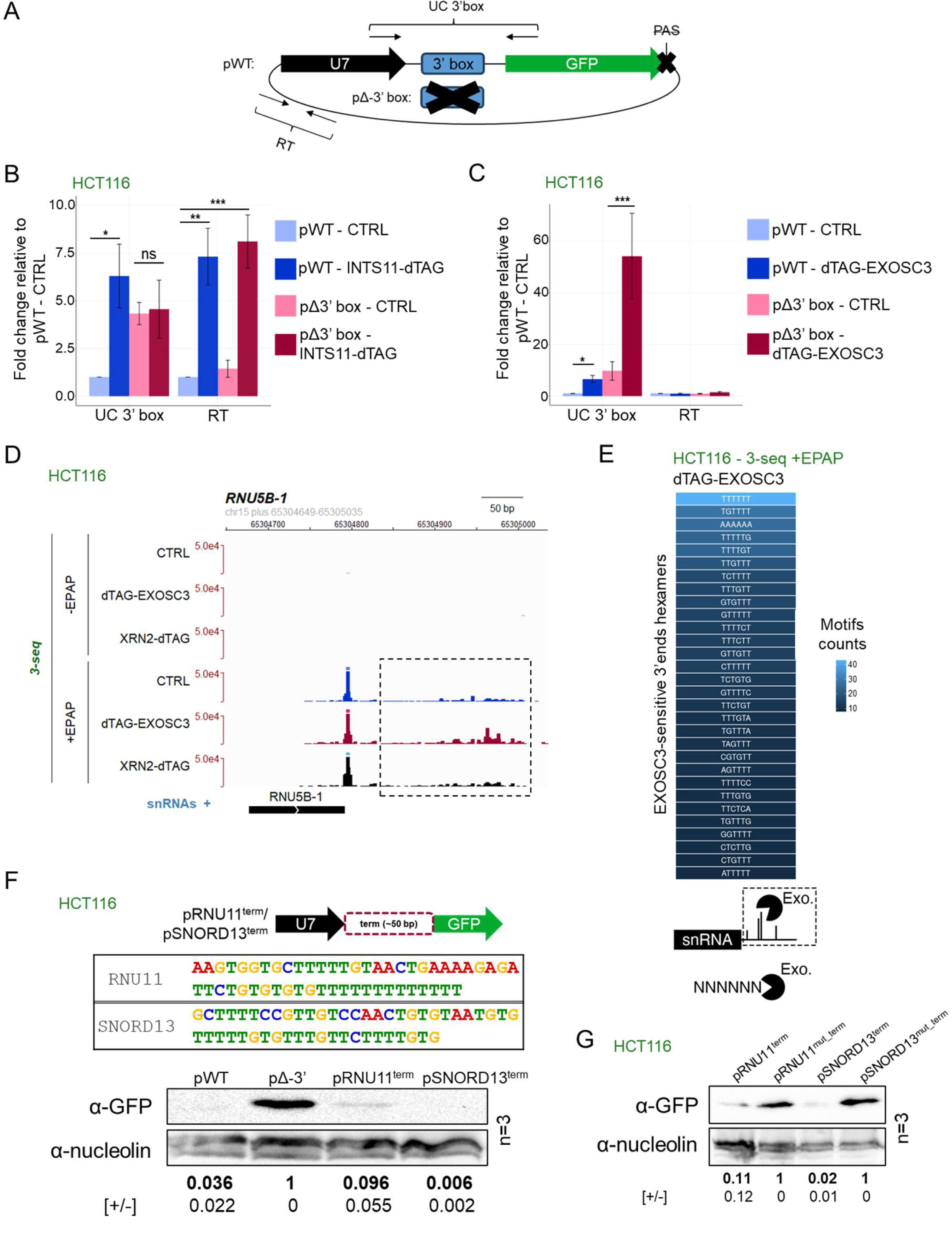
Termination of snRNA transcription is 3’ box-independent. A. Schematic representation of the pWT and pΔ3’ box reporter plasmids. The U7 snRNA sequence is followed by its 3’ box (or its deletion) and a GFP gene with a mutated PAS. Arrow pairs denote the two amplicons (‘UC 3’ box’ and ‘RT’) used for qRT-PCR analysis. B. qRT-PCR analysis of pWT or pΔ3’box reporter expression from *INTS11-dTAG* HCT116 cells untreated (CTRL) or treated (INTS11-dTAG) for 4hr with dTAGv-1. RNA spanning the 3’ box or from the downstream region was measured by ‘UC 3’ box’ and ‘RT’ amplicons, respectively. Measured RNA quantities were normalised to levels of GAPDH RNA. Mean fold-change values were calculated by comparative quantitation *vs.* pWT CTRL. n=4. Error bars = standard error of the mean (SEM). *, ** and *** denote p≤.05, .01 and .001. C. As in (B) but performed in *dTAG-EXOSC3* HCT116 cells. n=4. Error bars = SEM. *, and *** denote p≤.05 and .001. D. Genome browser view of RNA 3’-seq signal over the *RNU5B-1* locus and deriving from +/− EPAP treated RNA from HCT116 (CTRL), *dTAG-EXOSC3* or *XRN2-dTAG* cells treated with dTAGv-1 for 4hrs. The dashed box highlights a region with EXOSC3-sensitive 3’ ends. E. Sequence composition of the most common hexameric motifs at the 3’ terminal of 3’-seq reads (+EPAP) found ≤4kb downstream of snRNA TESs and stabilised by log_2_ change ≥1 following EXOSC3 depletion from dTAG-EXOSC3 cells *vs*. HCT116 CTRL cells (see below schematics). Note that motifs are shown as the coding DNA strand equivalent. The heat scale shows the motif count. F. Top panel: Schematic representation of utilized reporter construct and the sequences of the inserted *RNU11* and *SNORD13* downstream elements. Bottom panel: Western blotting analysis of GFP expression following transfection of HCT116 cells with the pWT, pΔ3’box, p^term^RNU11, or p^term^SNORD13 containing reporter constructs. Quantifications below the western membrane show relative GFP expression levels relative to those of the pΔ3’ box reporter. n=3 (+/− values = SEM) following normalisation to nucleolin protein levels. G. Western blotting analysis as in (F), but of the p^term^RNU11 and p^term^SNORD13 constructs and their mutated derivatives, in which all T’s were substituted for A’s. Quantitation of the GFP signal is relative to the respective mutant following normalisation to nucleolin protein levels. n=3 (+/− values = SEM).

Our results from Figure 1 showed that the exosome degrades the products of an INT-independent snRNA transcriptional termination process. This collectively predicted that snRNA termination will continue to generate exosome substrates even when a 3’ box is absent. To test this, we transfected the pWT and pΔ3’ box plasmids into *dTAG-EXOSC3* HCT116 cells, allowing rapid exosome inactivation (Figure S3A). Unlike the INTS11 depletion, which only affected UC 3’ box amplicon levels when the 3’ box was present, exosome depletion substantially enhanced the amount of RNA spanning the 3’ box region for both pWT and pΔ3’ box constructs (Figure 3C, ‘UC 3’ box’ columns). In contrast, exosome depletion did not affect RT RNA expression from either construct, presumably because the 3’ box-independent termination process remained fully operational and because the exosome has no direct effect on the transcription attenuation process. Therefore, a 3’ box-independent termination mechanism generates snRNA precursors, which are substrates for the exosome. We suggest that co-transcriptional 3’ box processing by the INT complex insulates the mature snRNA from this degradation initiating from downstream termination sites. Supporting this notion, published photoactivatable-ribonucleoside-enhanced cross-linking and immunoprecipitation (PAR-CLIP) data of the exosome ribonuclease DIS3 (Szczepinska et al. 2015) demonstrated its elevated binding to regions downstream of the snRNA 3’ box (Figure S3B).

If the INT complex is not terminating snRNA transcription, then what is? We previously detected signs of snRNA transcription termination at T-rich elements (Davidson et al. 2020). To explore this systematically, we generated further RNA 3’-seq data to map the 3’ ends of newly produced transcripts but employing dTAG degron HCT116 cells for rapid EXOSC3 depletion or, to test any contribution of 5’→3’ degradation, XRN2 (Figure S3A). As expected, the *RNU5B-1* locus showed no read coverage in the -EPAP conditions (Figure 3D, top panels). In contrast, +EPAP samples displayed the products of 3’ box cleavage (Figure 3D, +EPAP panels). Moreover, numerous downstream 3’ ends were revealed and were stabilised by EXOSC3 loss. This finding is consistent with results from Figure 1C, showing reproducibility between data obtained following siRNA-mediated depletion of EXOSC3 from HeLa cells and degron-depletion of EXOSC3 from HCT116 cells. XRN2 depletion did not impact positions or levels of downstream 3’ends in line with its inability to terminate snRNA transcription (Eaton et al. 2018).

To assess the contribution of DNA/RNA sequence to snRNA transcription termination in an unbiased manner, we extracted the most common 3’ terminal hexamers that were upregulated by log_2_ change ≥1 in +EPAP 3’-seq samples from dTAG-treated dTAG-EXOSC3 *vs.* CTRL HCT116 cells. We surmised that these exosome-degraded transcripts were terminated RNAs as indicated by our experiments in Figure 1 and that nucleotides immediately upstream of the exosome-sensitive 3’ ends should represent any involved DNA sequence motifs. To be conservative, we searched for hexameric sequences knowing that T≥4 is sufficient to induce DNA-directed transcriptional termination of RNAPIII (Arimbasseri et al. 2013). Plotting sequences as their DNA coding strand equivalent revealed T6 as the most enriched hexamer and several additional T-rich elements (Figure 3E). Importantly, this did not simply reflect a T bias of the underlying DNA strand because the percentage of exosome-sensitive 3’ ends terminating at coding strand Ts exceeded the percentage of Ts in the same genomic sequence (Figure S3C).

To test whether naturally occurring T-rich elements could act as terminators when transplanted into reporter DNA, we replaced the 3’ box within our U7-GFP reporter (retaining the GFP PAS in this case) with ∼50 nucleotides of T-rich elements from downstream of *RNU11* or *SNORD13* sn/snoRNA TESs. These contained visible clusters of EXOSC3-sensitive 3’ ends (Figure S3D). The resulting plasmids were transfected into HCT116 cells along with the pWT and pΔ3’ box reporter controls and GFP expression was assayed by western blotting analysis. As expected, GFP levels were low for pWT- and high for pΔ3’ box constructs, respectively (Figure 3F). Replacing the 3’ box with *SNORD13* or *RNU11* sequence strongly reduced GFP expression, suggesting these sequences terminate transcription. We then substituted all Ts within the cloned elements with A’s, which resulted in the full recovery of GFP expression (Figure 3G). This strongly implied T-specific termination and that A-rich elements do not harbour the same capacity. We conclude that transcriptional termination of snRNAs (and independently transcribed snoRNAs) coincides with T-rich elements and yields RNA substrates for the exosome.

### Genome-wide transcriptional termination at T-rich elements

Having established snRNA transcription termination at T-rich elements, we wondered whether such termination might be more widespread. Beyond pre-snRNAs, many transcripts are exosome-sensitive, which might in part be explained by termination over T-stretches. Hence, we broadly assessed the exosome-sensitivity of newly synthesised RNA, reasoning that this would predict such DNA-directed termination. We used TT-seq from siCTRL-or siEXOSC3-treated HeLa cells (Lykke-Andersen et al. 2021), and plotted the log_2_ change in signal for siEXOSC3 *vs*. siCTRL samples over the bodies of all TUs with TT-seq signal detected downstream of their annotated TESs. This analysis revealed the expected production of exosome-sensitive transcripts (Figure 4A). We previously demonstrated that RNAs deriving from mono-exonic TUs are more exosome-sensitivity than RNAs from multi-exon TUs (Lykke-Andersen et al. 2021). Indeed, stratifying the TT-seq analysis by mono- and multi-exon TUs recapitulated this finding (Figure 4B). The production of unstable RNA from gene bodies and downstream regions of mono-exonic TUs was expected because transcription terminates within the ∼3kb broad promoter-proximal region by a combination of mechanisms (Austenaa et al. 2015; Austenaa et al. 2021; Estell et al. 2021; Estell et al. 2023; Rouviere et al. 2023). Consistent with this, exosome sensitivity was also observed within TSS-proximal regions of multi-exon TUs; however, there was significantly less exosome-sensitive RNA deriving from the gene bodies of this TU class (Figure 4B). Finally, exosome-sensitive RNA production was also detectable downstream of some multi-exon TESs. This was somewhat surprising because RNA downstream of the PAS was expected to be largely degraded by XRN2 in a process linked to PAS cleavage (Rodriguez-Molina et al. 2023). We conclude that exosome-sensitive RNAs are abundant within TSS- and, to a lesser extent, TES-proximal regions.

**FIGURE 4:**
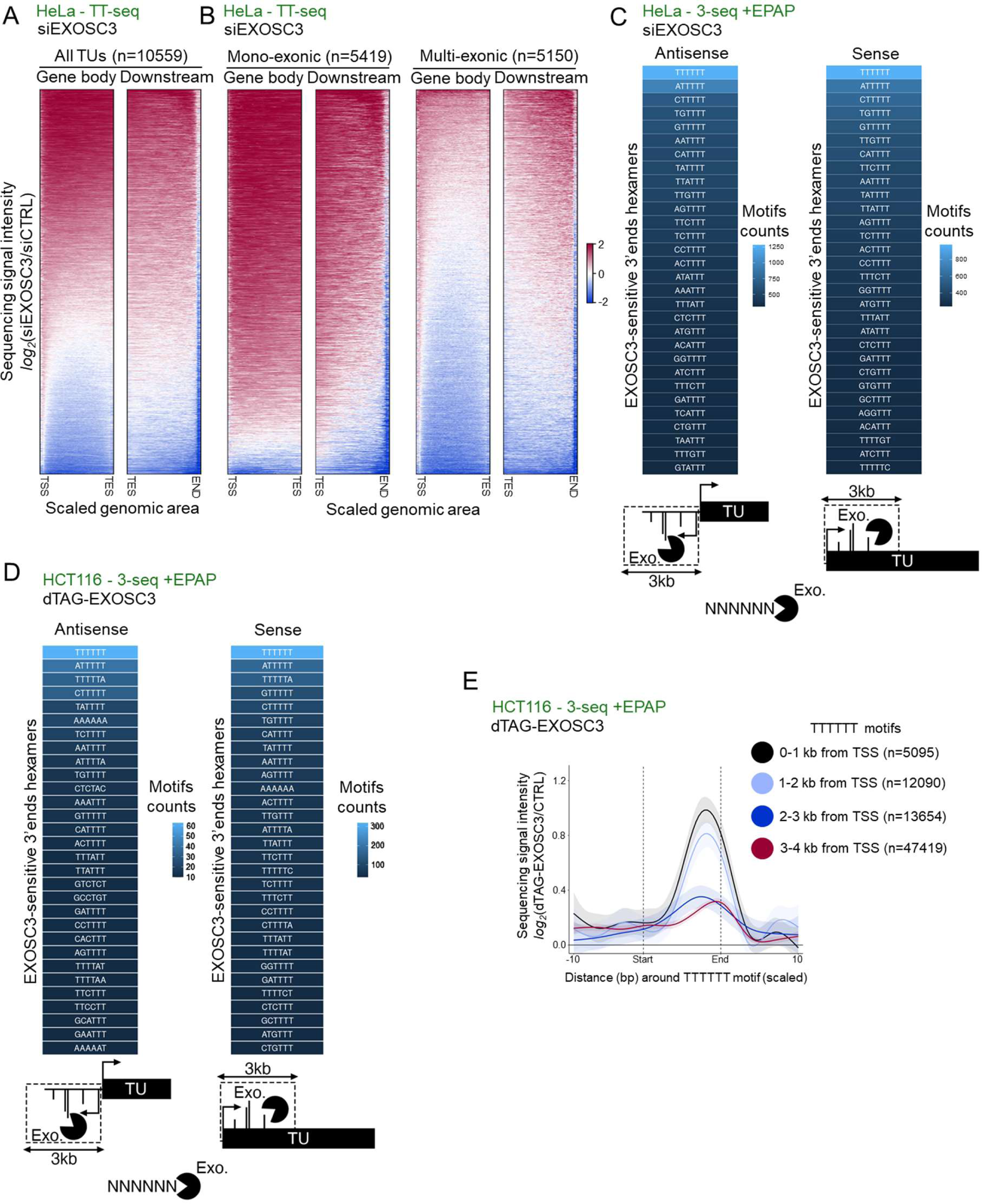
Global transcriptional termination at T-rich elements. A. Heatmaps showing TT-seq data from siCTRL- and siEXOSC3-treated HeLa cell RNA samples (GSE151919 (Lykke-Andersen et al. 2021)). The left-hand heatmap shows the log_2_ change in sequencing signal intensity (siEXOSC3 *vs.* siCTRL) over the gene body region (TSS to TES) of all TUs (n=10559) and the right-hand heatmap shows the same analysis but for the area downstream of the TES. B. Heatmaps as in (A) but showing the gene body and TES-downstream regions of mono-exonic (left-hand two maps) and multi-exonic (right-hand two maps) TUs. C. Sequence composition of the most common hexameric motifs immediately upstream of the 3’ end of 3’-seq (+EPAP) reads over regions within 3kb sense or antisense of TSSs and stabilised by log_2_ change ≥1 following siEXOSC3 *vs.* siCTRL treatment of HeLa cells. Note that motifs are shown as the coding DNA strand sequence. The heat scale shows motif counts. Analyses derives from (GSE151919 (Lykke-Andersen et al. 2021)). D. Analysis as in (C), but in dTAG-treated *dTAG-EXOSC3* HCT116 cells *vs.* CTRL HCT116 cells. E. Meta-analysis of +EPAP 3’-seq coverage over ≥T6 regions within 0-1, 1-2, 2-3, or 3-4kb from multi-exon TU TSS’s. The x-axis shows this region (start-end), including 10 nucleotides upstream and downstream of it. The y-axis displays the log_2_ change in signal sequence intensity in dTAG-treated *dTAG-EXOSC3* or *XRN2-dTAG* HCT116 cells *vs.* HCT116 CTRL cells.

Given our observations at snRNA loci (Figure 3E), we hypothesised that transcription termination at T-rich elements might also account for at least some exosome-sensitive 3’ ends within mono- and multi-exon loci. Accordingly, we used our RNA 3’-seq data to identify the most common hexameric motifs immediately upstream of exosome-sensitive RNA 3’ ends within 3kb upstream antisense and downstream sense of protein-coding gene TSSs. In both cases, T6 was the most common motif in HeLa (Figure 4C) and HCT116 (Figure 4D) cells. Again, this did not simply reflect a T bias within the underlying DNA coding strand because the percentage of exosome-sensitive 3’ ends terminating at coding strand Ts exceeded the percentage of Ts found in the genome at these regions (Figure S4A and S4B). We propose that many promoter-proximal exosome substrates are generated by transcriptional termination at T-tracts/T-rich elements.

Previous data imply that, after transcription initiation, full elongation competence is gradually established as RNAPII accelerates throughout a ∼3kb “pausing zone” (Jonkers et al. 2014; Fong et al. 2022). Suggestive of this region involving DNA-directed termination, exosome-sensitive T6 termini were most frequent within the first 1kb proximal to the TSS whereafter prevalence decreased with increasing distance to the promoter (Figure 4E). An example of such promoter-proximal termination over an exosome-sensitive T-tract is provided in Figure S4C. Thus, promoter-proximal RNAPII is most sensitive to transcriptional termination over T-rich sequences and its susceptibility to this mechanism diminishes as it moves into the gene. This implied that mature RNAPII elongation complexes do not terminate at T-tracts. To generally analyse termination at T-tracts across multi-exon TUs, we assayed the exosome-or XRN2-sensitivity of T≥6 termini over different regions of the gene body and downstream of the TES. Consistent with the data from Figures 4C and 4D, there was substantial exosome sensitivity at T-tracts in the promoter-proximal region (Figure S4D, left panel). A limited level of exosome-sensitive at T-tracts was also observed downstream of the TES (Figure S4D, right panel). There was also very little exosome-sensitivity at T≥6 motifs over the gene body, even though more such motifs were present because some genes are long (Figure S4D, mid panels). XRN2 loss caused no accumulation of RNA terminating over T-tracts over any of these regions. These data demonstrate that termination at T-rich elements is the most prevalent promoter-proximally. However, mature gene body elongation complexes are much less prone to terminate at T-rich elements.

### DNA-directed and torpedo termination mechanisms are largely independent

XRN2-mediated transcriptional termination is the major mechanism downstream of the PAS. In contrast, DNA-directed termination appears to be most common in promoter-proximal regions. These observations might reflect the two transcriptional termination mechanisms operating in spatially separate contexts. We tested this hypothesis by comparing the exosome- and XRN2-sensitivity of transcripts from within and downstream of mono- and multi-exon TUs. To do so, we generated nuclear RNA-seq data from *dTAG-EXOSC3* HCT116 cells and compared it to our previously published nuclear RNA-seq generated after the rapid loss of XRN2-mAID (depleted *via* the auxin-inducible degron (Eaton et al. 2018)) from HCT116 cells. Consistent with the RNAi-induced depletion of EXOSC3 in HeLa cells (Figure 4B), rapid EXOSC3 elimination from HCT116 cells revealed that mono-exonic RNAs and promoter-proximal transcripts from multi-exon TUs were the most exosome-sensitive with a milder effect evident downstream of a subset of multi-exon TESs (Figure 5A). In contrast, mono-exonic RNAs and promoter-proximal transcripts from multi-exon TUs were largely unaffected by XRN2 loss. Instead, and as expected due to the widespread torpedo mechanism, RNA deriving from downstream of multi-exon TESs was near-universally stabilised by XRN2 loss. Hence, DNA-directed termination is most prevalent promoter-proximally whereas Torpedo termination is the dominant mechanism downstream of the PAS.

**FIGURE 5:**
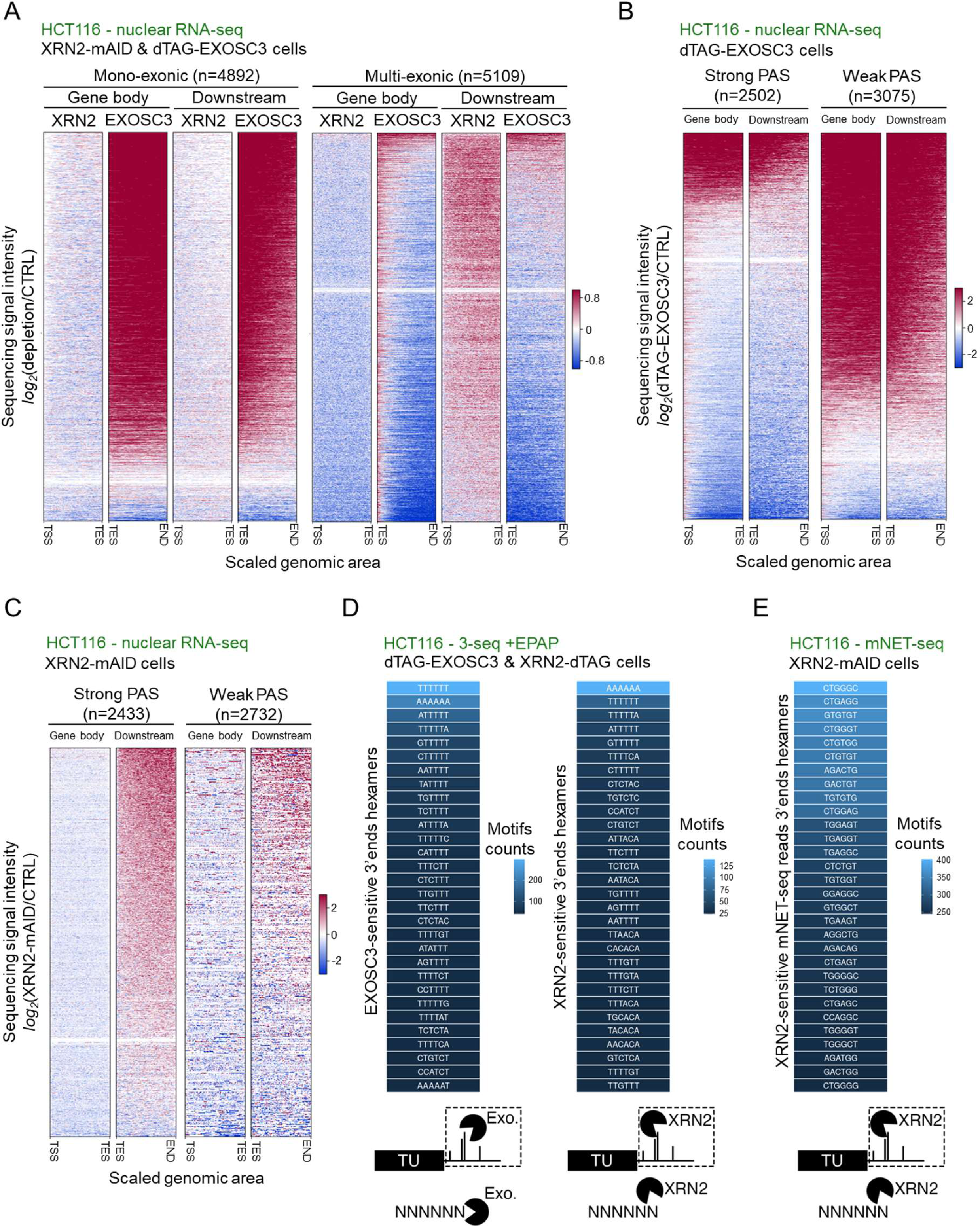
Exosome and XRN2 activities denote separate use of DNA-directed and torpedo termination. A. Heatmap analysis as in Figure 4A, but showing the log_2_ change in nuclear RNA-seq signal intensity between samples obtained after the rapid loss of XRN2 from XRN2-mAID HCT116 cells (XRN2 – data from GSE109003 (Eaton et al. 2018)) or EXOSC3 from *dTAG-EXOSC3* HCT116 cells (EXOSC3) *vs.* their respective controls. The left-hand four heatmaps show gene body and downstream regions of mono-exonic TUs and the right-hand equivalents display the same regions from multi-exon TUs. B. Heatmap analysis as in (A) but showing log_2_ change in nuclear RNA-seq signal intensity between -dTAG-EXOSC3 *vs.* CTRL samples over the gene body- and downstream-regions of TUs with predicted strong (left two heatmaps) or weak (right two heatmaps) PASs. C. As (B) but for XRN2-mAID HCT116 cells depleted of XRN2 (Eaton et al. 2018). D. Sequence composition analysis as in Figure 3E, but of 3’ ends derived from downstream of multi-exonic TU TESs and stabilised by log_2_ change ≥1 following dTAG depletion of EXOSC3 or XRN2 *vs*. their HCT116 cell CTRL. E. Sequence composition analysis as in (D), but of 3’ ends of mNET-seq reads stabilised by log_2_ change ≥1 following XRN2-mAID depletion *vs*. untreated CTRL HCT116 cells. The data are from GSE109003 (Eaton et al. 2018).

While RNA cleavage, usually at a PAS, is obligatory for the torpedo termination, it is conceptually unnecessary for a DNA-directed process. We therefore hypothesised that the efficiency of PAS cleavage might dictate the use of either termination pathway downstream of multi-exon gene TESs. To examine this, we used APPARENT2 – a machine learning model trained to predict the probability of PAS cleavage for all 23145 expressed HeLa transcripts (see Materials and Methods). These were then subdivided into two classes, representing those most and least likely to undergo cleavage (assigned as ‘strong’ or ‘weak’ PAS). As expected, protein-coding transcripts were the most enriched biotype with strong PASs, whereas weak signals were primarily assigned to ncRNAs less likely to undergo PAS-dependent processing (Figure S5A) (Schlackow et al. 2017; Davidson et al. 2019; Lykke-Andersen et al. 2021). To demonstrate the biological validity of these *in silico* predictions, we analysed our published nuclear RNA-seq data obtained after the auxin-induced depletion of CPSF30 (Estell et al. 2021). Because CPSF30 is essential for PAS recognition and consequent transcriptional termination (Clerici et al. 2017; Sun et al. 2018), its depletion was predicted to increase RNA signals beyond all genuinely processed PASs. This was indeed the case for the strong PAS predictions, whereas the predicted weak PASs were mostly unaffected by CPSF30 depletion (Figure S5B).

We then assessed the effect of EXOSC3 or XRN2 loss on nuclear RNA signals upstream and downstream of weak or strong PASs. Exosome sensitivity was most prevalent upstream and downstream of the predicted weak PASs, reflecting its known ability to degrade ncRNAs to completion (Figure 5B). In contrast, XRN2 sensitivity was strongest downstream of efficiently cleaved PASs and its depletion did not impact RNA levels over gene bodies (Figure 5C). Although we only had XRN2-depletion data available from HCT116 cells, TT-seq data deriving from siEXOSC3-treated HeLa cells gave a result similar to the dTAG depletion of EXOSC3 from HCT116 cells (Figure S5C). Further leveraging our comprehensive TU annotation in HeLa cells (Lykke-Andersen et al. 2021), we identified T6 as the most common six nucleotide motif at 3’ ends stabilised by EXOSC3 loss downstream of weak PASs (Figure S5D). Although T6 was also the most enriched motif downstream of strong cleavage sites, its abundance was ∼10-fold lower than for weak PASs (Figure S5D, note difference in motifs counts scaling) in line with DNA-directed termination being less frequent here. Taken together, these analyses reveal that exosome and XRN2 activities are usually employed in different locations/circumstances. Specifically, XRN2-dependent termination is associated with PAS cleavage, which is unnecessary for the DNA-directed process.

DNA-encoded information could still contribute to the torpedo mechanism through a scenario where XRN2 and the exosome cooperatively degrade the fragment downstream of PAS cleavage. If so, XRN2 depletion might also enrich for specific motifs at the terminus of 3’-seq reads downstream of multi-exon TUs and we therefore compared these over such regions upon EXOSC3-or XRN2-loss in their respective HCT116 degron cells. Consistent with previous analyses, T6 was the top hit after EXOSC3 depletion with other T-rich regions also enriched (Figure 5D, left panel). Following XRN2 loss, A6 was the most common 3’ motif, which should be interpreted cautiously because of the employed EPAP activity (Figure 5D, right panel). However, although T6 was second-ranked, it was recovered at a ∼4-fold lower frequency than upon EXOSC3-depletion, despite the RNA downstream of the PAS being generally more stabilised upon XRN2-depletion. The recovery of XRN2-targeted transcripts with T6 3’ termini might reflect some cooperation between DNA-directed and Torpedo termination or the post-transcriptional degradation of RNA already released by DNA-directed termination.

The torpedo mechanism occurs *via* a “sitting duck” mechanism, whereby RNAPII slows down beyond the PAS and awaits termination by XRN2 (Cortazar et al. 2019). If T-tracts enable XRN2-dependent termination, these “sitting duck” RNAPIIs might be found over such sequences. If so, XRN2 depletion would trap chromatin-associated RNAPII over T-tracts. To test this, we reanalysed our published mammalian native elongating transcript (mNET)-seq data obtained after the rapid auxin-induced loss of XRN2 in XRN2-mAID cells (Eaton et al. 2018). mNET-seq maps the exact position of RNAPII by sequencing the 3’ end of the RNA residing in its active site (Nojima et al. 2015). We extracted the most frequent XRN2-sensitive 3’ terminal hexamer motifs at the 3’ end of mNET-seq reads downstream of the PAS. Unlike 3’ ends stabilised by EXOSC3 loss, and visualised by 3’-seq, these ends were G-rich and T6 was not among the most abundant motifs (Figure 5E). This is consistent with previous observations that G-rich elements can act as RNAPII pause sites and facilitate XRN2-dependent termination (Gromak et al. 2006; Sheridan et al. 2019; Fong et al. 2022). Based on this finding, we suggest that RNAPII does not frequently pause/arrest over T-rich elements before XRN2-dependent termination.

### Selected short protein-coding genes employ both DNA-directed and Torpedo termination

There has been a long debate concerning the necessity of PAS cleavage for transcriptional termination downstream of protein-coding genes (Libri 2015). Multiple studies have demonstrated PAS-dependent XRN2-mediated termination *in vivo* (Fong et al. 2015; Eaton et al. 2018). Still, others – especially those conducted *in vitro* – have argued that transcription can also terminate independently of PAS cleavage (Nag et al. 2006; Zhang et al. 2015). DNA-directed termination that does not require RNA cleavage might help resolve this debate. To isolate protein-coding genes that may undergo DNA-directed transcriptional termination, we used nuclear RNA-seq data to select those whose 3’ flanking RNA was exosome-sensitive. With inclusion criteria of log_2_ change ≥1.5 upon dTAG-EXOSC3 loss from HCT116 cells and without any corresponding increase in the gene body, 43 such TUs were selected (Figure 6A). The latter criterion was important because RNAs from most exosome-sensitive TUs display gene body effects (e.g. Figure 5A). With only one exception, these 43 TUs were short (∼2.5kb on average), and most contained a canonical AAUAAA PAS hexamer (Figure S6A). Although their short lengths were more reminiscent of mono-exonic TUs, producing primarily exosome-sensitive RNAs, a low level of torpedo termination was evident based on increased PAS-downstream POINT-seq signal following XRN2 depletion (Figure 6B, Figure S6B).

**FIGURE 6:**
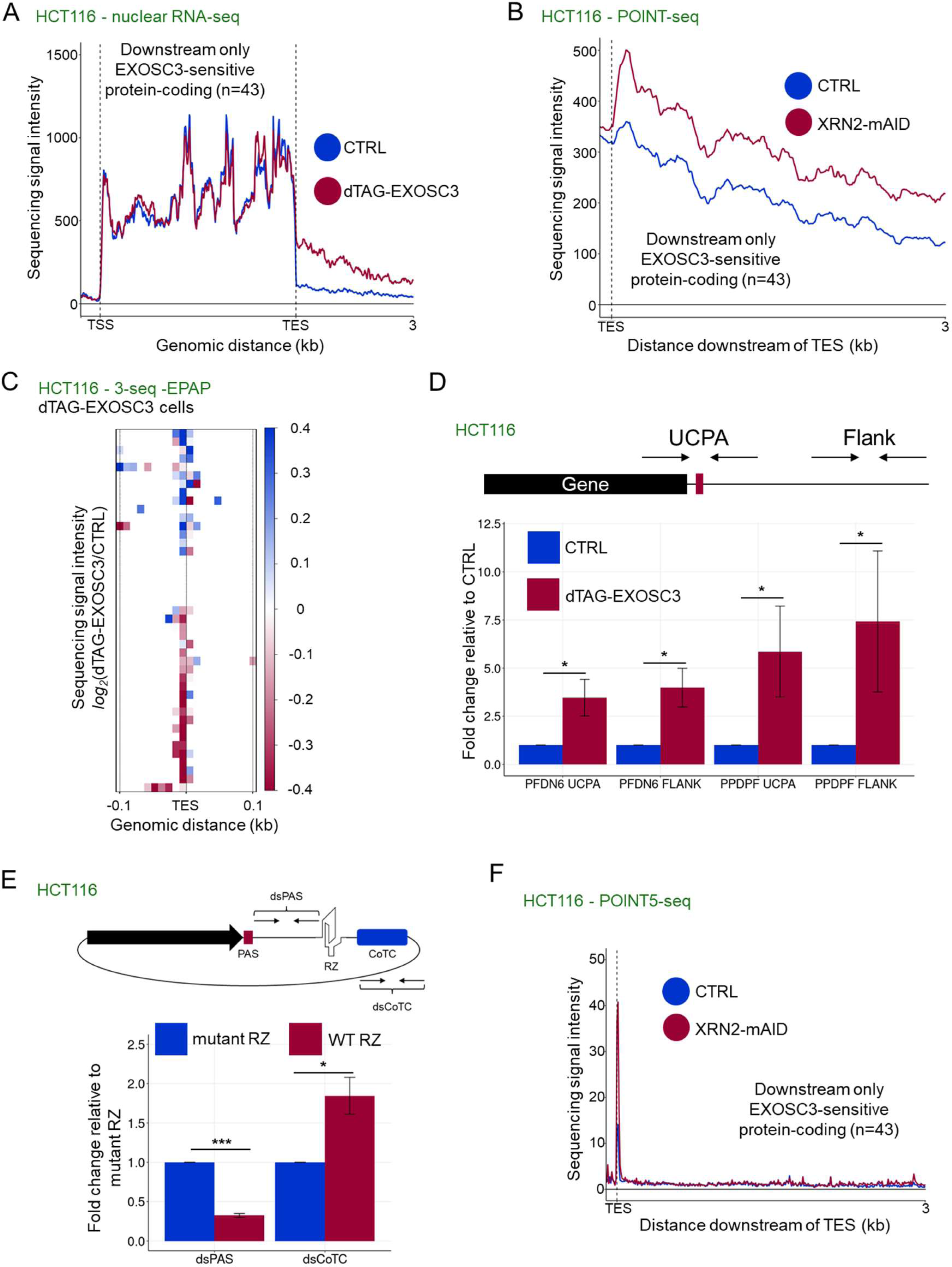
Mutually exclusive use of DNA-directed and torpedo termination downstream of some short protein-coding genes. A. Metaplot of nuclear RNA-seq signal from *dTAG-EXOSC3* HCT116 cells treated, or not, for 4hr with dTAGv-1. The plot displays signals over 43 protein-coding TUs with a log_2_ change ≥1.5 increase in signal intensity downstream of the TES following EXOSC3 depletion. The x-axis shows the gene body region (TSS-TES) and 3kb of the downstream region. The y-axis sequence signal intensity units are RPKM. B. Metaplot as in (A) but for POINT-seq signal derived from *XRN2-mAID* cells treated, or not, for 4hr with auxin (GSE159326 (Sousa-Luis et al. 2021)). The y-axis sequence signal intensity units are BPM. C. Heatmap showing the log_2_ change in sequencing signal intensity in -EPAP 3’-seq signal samples (dTAG-treated *dTAG-EXOSC3* HCT116 cells *vs*. CTRL HCT116 cells). The x-axis is centered around the TESs of the 43 TUs analysed in (A) and displays 0.1kb of respective upstream and downstream regions. D. qRT-PCR analysis of non-PAS cleaved (‘UCPA’ amplicon) and 3’ flanking (‘Flank’ amplicon) RNAs from the *PFDN6* and *PPDPF* genes. Samples were from dTAG-EXOSC3 cells treated, or not, with dTAGv-1 for 4hrs (dTAG-EXOSC3 and CTRL, respectively). Measured RNA quantities were normalised to GAPDH RNA levels. Mean fold-change values were calculated by comparative quantitation and plotted relative to those obtained in CTRL conditions. n=3. Error bars = SEM. * denotes p≤0.05. E. qRT-PCR analysis of RNA derived from human β-globin plasmids, containing a hepatitis δ-ribozyme (‘WT RZ’), or its inactive mutant (‘mutant RZ’) inserted between the PAS and CoTC elements as indicated. Employed amplicons were positioned between the PAS and the CoTC (‘dsPAS’) and downstream of the CoTC (‘dsCoTC’), respectively. Measured RNA quantities were normalised to GAPDH RNA levels. Mean fold-change values were calculated by comparative quantitation and plotted relative to those obtained in CTRL conditions. Error bars = SEM. n=4. * and *** denotes p≤0.05 and p≤0.001. F. Metaplot of published POINT5-seq data from XRN2-mAID *vs.* CTRL cells (GSE159326 (Sousa-Luis et al. 2021)). The 43 TUs from (A) are shown. The x-axis shows the TES and 3kb of downstream sequence. The y-axis sequence signal intensity units are BPM. The only visible XRN2-sensitive 5’ end coincides with the TES (corresponding to PAS cleavage).

By seemingly employing both interrogated termination mechanisms, these 43 TUs were appropriate to dissect such inter-/co-dependences. Therefore, we asked whether exosome degradation of 3’ flanking RNA from these TUs was compatible with PAS cleavage, which would be necessary for DNA-directed termination to cooperate with the torpedo mechanism. To do this, we turned to our RNA 3’-seq data obtained without EPAP treatment (to detect naturally cleaved and polyadenylated mRNA). If exosome degradation from a region downstream of the PAS competes with PAS cleavage, we would expect more processed mRNA upon exosome loss. However, EXOSC3-depletion only modestly affected 3’ end processing without a conclusive outcome (Figure 6C). Furthermore, qRT-PCR analysis of two example transcripts (PFDN6 and PPDPF), with primer pairs spanning their respective PASs (‘UCPA’) or 3’ flanks (‘Flank’), showed significant stabilisation of unprocessed RNA upon exosome loss (Figure 6D). Thus, exosome activity appears most relevant for transcripts that do not undergo PAS cleavage.

The dual EXOSC3- and XRN2-sensitivities of these transcripts and their short TU lengths were reminiscent of our original studies on transcriptional termination, using the similarly short (1.8 kb) human β-globin gene as a model (Dye and Proudfoot 2001; West et al. 2004; West et al. 2006). Here, transcription termination requires a PAS, and a downstream termination sequence, termed the Co-Transcriptional Cleavage (CoTC) element. We originally proposed that RNA transcribed from this terminator was co-transcriptionally cleaved based on its apparent discontinuity and on the dual degradation of the resulting RNA fragment by both the exosome and XRN2 (Dye and Proudfoot 2001; West et al. 2004; West et al. 2006). However, the identity of any CoTC endonuclease remained a mystery. Inspired by our findings here, we hypothesised that DNA-directed termination at the CoTC element, rather than RNA cleavage, would generate an exosome substrate. In contrast, the XRN2 entry site would be generated exclusively by PAS cleavage. To test this possibility, we generated a β-globin reporter with a hepatitis δ ribozyme (RZ), or an inactive mutant RZ, inserted between the PAS and CoTC elements (Figure 6E, top panel). RZ cleavage yields an RNAPII-associated cleavage product with an XRN2-resistant 5’OH end, thereby inhibiting the torpedo termination (Stevens and Maupin 1987; Eaton et al. 2020). Hence, if XRN2 degradation initiates solely from the cleaved PAS, RZ cleavage would block its progress and stabilise the downstream RNA. However, if a downstream CoTC provides an additional XRN2 entry site, the downstream RNA would not accumulate. These constructs were transfected into HCT116 cells and qRT-PCR was used to assay upstream and downstream RNA levels. The RZ sequence increased read-through transcription beyond the CoTC element compared to its mutated RZ counterpart (dsCoTC amplicon, Figure 6E), suggesting that the CoTC element cannot generate an XRN2 substrate. The upstream RZ cleavage product (dsPAS amplicon) can be degraded (Muniz et al. 2015), explaining the lower level of this species for the WT *vs.* the mutant RZ. These data are compatible with CoTC being a DNA-directed terminator.

To test whether PAS cleavages provide the only detectable XRN2 entry sites of the RNAs derived from the 43 TUs, displaying strong exosome sensitivity over their 3’ flanks, we analysed published POINT5-seq data performed after XRN2 depletion (Figure 6F). Meta-analysis of the 3’ flanking regions showed a single XRN2-sensitive peak, corresponding to the PAS cleavage position. This further suggested that the exosome-sensitivity of these RNAs derives directly from termination rather than from the endonucleolytic cleavage of the 3’ flanking RNA. Although our 3’-seq data had low coverage downstream of these genes, some exosome-sensitive T-rich termini were recovered (Figure S6C). Thus, the DNA-directed and torpedo termination are used mutually exclusively at the protein-coding genes that employ both mechanisms. Because DNA-directed termination does not require PAS cleavage, it can be utilised if 3’ end processing fails and when the XRN2 torpedo cannot engage the RNAPII-associated RNA.

Finally, RDH transcripts represent an exceptional class of protein-coding RNAs because their 3’ ends are non-polyadenylated even though they undergo 3’ end formation using the CPSF73 endonuclease (Marzluff and Koreski 2017). However, like the 43 TUs analysed here, they are short (<1kb) and XRN2 has a more limited impact on their transcriptional termination (Fong et al. 2015; Eaton et al. 2018). Thus, DNA-directed termination might play a role here. Consistent with this notion, RDH transcripts displayed robust exosome sensitivity downstream of their TES (Figure S6D). Moreover, T-rich hexamers upstream of EXOSC3-targeted 3’ ends downstream of RDH TESs were enriched, with more such 3’ ends terminating at Ts *vs.* the percentage of genomic Ts, in both HCT116 (Figures S6E and S6F) and HeLa (Figures S6G and S6H) cells. We conclude that a range of short human protein-coding genes, including RDH TUs, employ DNA-directed termination downstream of their TES.

## DISCUSSION

The RNA signals required for PAS processing and their connection to transcriptional termination were defined decades ago (Proudfoot 2011). More recently, the ability of the INT complex to elicit RNA cleavage-dependent transcriptional attenuation was also revealed (Elrod et al. 2019; Tatomer et al. 2019; Lykke-Andersen et al. 2021). Instead, prokaryotic RNAP and eukaryotic RNAPIII directly terminate at DNA elements, specifically at T-tracts. Here, we describe the widespread termination of mammalian RNAPII at T-rich elements (Figure 7). This mechanism is most relevant to RNAPII complexes located promoter-proximally and downstream of a subset of PASs. While we cannot exclude a function for unidentified factor(s) in promoting this termination, the importance of DNA sequence is underlined by the enrichment of T-stretches at exosome-sensitive 3’ termini in these regions and by the termination of reporter plasmid transcription by natural T-rich terminators.

**FIGURE 7:**
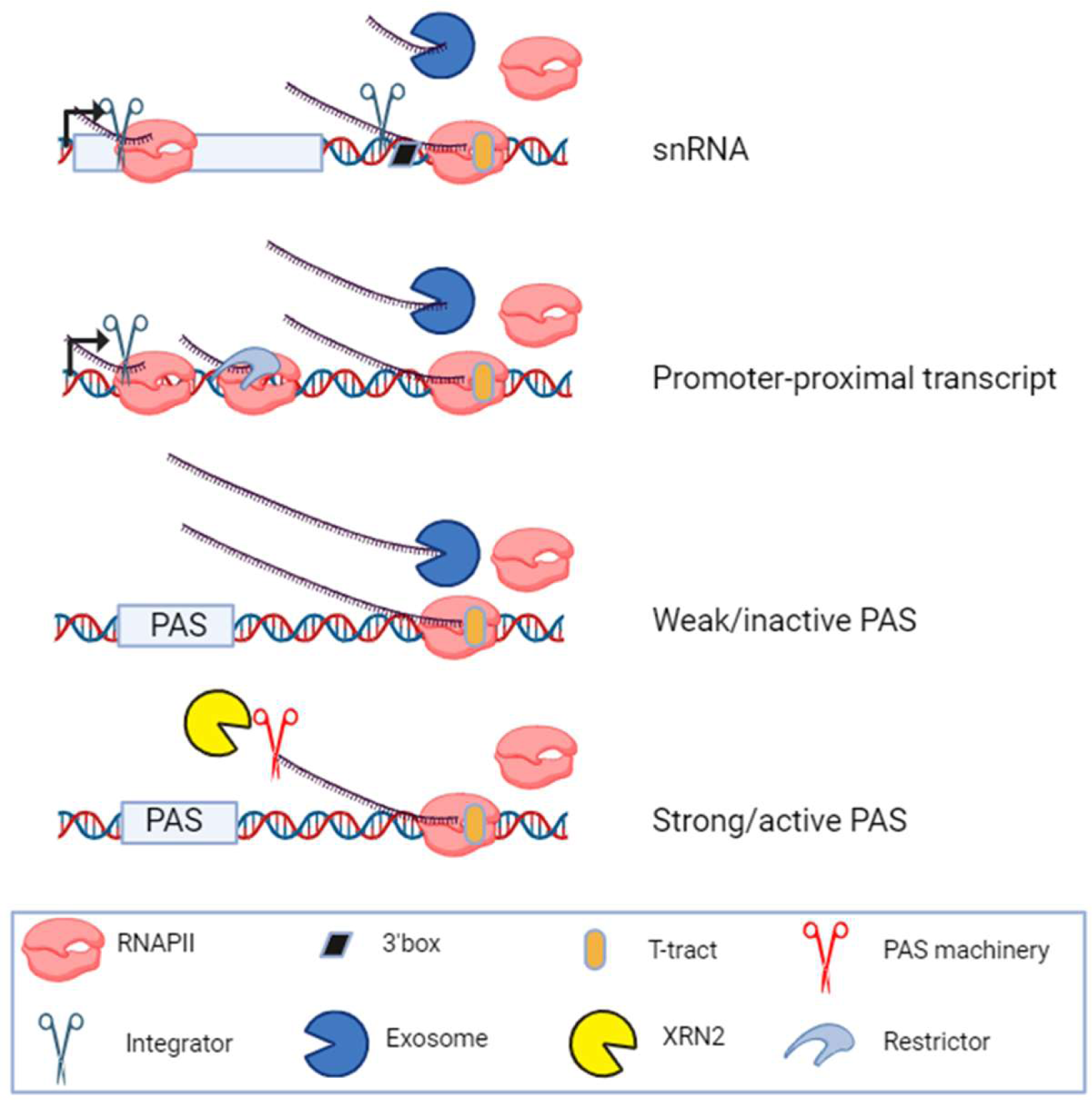
DNA-directed *vs*. torpedo-mediated transcription termination mechanisms. Top two schematics: At snRNA genes, co-transcriptional RNA 3’ box processing by the INT complex precedes but is not required for DNA-directed termination and subsequent exosome degradation. INT also attenuates snRNA transcription, explaining the apparent transcriptional read-through seen upon its depletion. In promoter-proximal regions, termination can occur *via* multiple complexes with INT and Restrictor being prominent examples. However, like for snRNAs, promoter-proximal termination can also occur by a DNA-directed process when RNAPII encounters T-rich elements in the coding strand of DNA. Lower two schematics: At the ends of multi-exon TUs (like protein-coding genes), efficient PAS cleavage leads to torpedo-directed transcription termination by XRN2 in most cases. Where PAS cleavage is inefficient or fails, DNA-directed termination provides a means of evicting RNAPII from the chromatin template. This might include cases where full elongation competence is not yet established (e.g. on some short protein-coding genes).

Although we did not analyse the DNA-directed termination of mammalian RNAPII in a purified system, several decades-old papers have addressed this, and the results support our findings. Purified mammalian RNAPII was shown to terminate over endogenous sequences and at bacterial termination elements without an apparent requirement for additional factors (Dedrick et al. 1987; Reines et al. 1987). When mapped, these termination sites indeed coincided with T-tracts or T-rich elements. A notable feature of such *in vitro* termination assays was that some T-tracts were ineffective RNAPII terminators and that RNAPII generally terminated less efficiently than prokaryotic RNAP over bacterial termination sequences. This mirrors what happens *in vivo* because RNAPII does not terminate at every T-tract it encounters. In contrast, mammalian RNAPIII termination is highly efficient at the first T≥4-tract in the DNA coding strand. In structures of terminating RNAPIII (Hou et al. 2021), poly-dT in the coding DNA strand orients into the polymerase exit tunnel where it is stabilised by a so-called fork loop. This is proposed to be critical for the DNA-directed termination process. In RNAPII structures (Hou et al. 2021), this exit tunnel is slightly broader possibly explaining why DNA-directed RNAPII termination is less efficient. An inefficient response to T-rich elements is presumably important to provide an opportunity to establish full RNAPII elongation competence.

We revealed that INT-dependent 3’ box cleavage serves a processing function at snRNAs, but that it is uncoupled from transcriptional termination, which instead occurs over T-tracts. This contrasts with the CPA complex at the PAS, which links mRNA 3’ end processing with transcriptional termination *via* the activity of XRN2 (Eaton et al. 2018). Initially appearing to contradict multiple observations of snRNA transcriptional read-through following INT depletion (Baillat et al. 2005; O’Reilly et al. 2014), which we also observed using different experimental approaches, we propose that this increased RNAPII activity is due to INT functioning to attenuate snRNA transcription. Alleviation of such attenuation, following INT depletion, increases snRNA transcription over the gene body and downstream region, which has been misinterpreted as a transcriptional termination defect. This explanation predicts INT loss to increase transcription before the 3’ box is encountered, which we observed using TT-seq and POINT-seq approaches and was evident in published Precision Run On-seq data (Stein et al. 2022). We suggest that INT has two roles at snRNA loci: attenuating their transcription and processing their resulting RNA 3’ box sequences. In other locations, INT presumably only performs its attenuation function since 3’ box consensus elements are inactive.

DNA-directed termination of budding yeast RNAPII was recently described to occur downstream of the PAS (Han et al. 2023). Here, we report its global use by mammalian RNAPII, which is important given the greater complexity of mammalian genes and genomes: they produce thousands of RNAs terminated in this manner, snRNA transcription terminates by this mechanism and regulated pausing of RNAPII is generally more prevalent in higher eukaryotes (Adelman and Lis 2012). In most cases, and exemplified at snRNA loci, DNA-directed termination works independently of RNA processing. Consequently, DNA-directed, and XRN2-dependent termination often work separately downstream of the PAS. When PAS cleavage occurs, transcriptional termination is XRN2-dependent; however, DNA-directed termination is utilised downstream of weak PASs or when PAS cleavage is absent. At times, XRN2 may terminate RNAPII positioned over T-tracts or could post-transcriptionally degrade RNA after DNA-directed termination. Indeed, the products of termination over T-tracts accumulate following Rat1 depletion from budding yeast, and T-tracts facilitate Rat1-dependent termination *in vitro* (Han et al. 2023). Supporting our hypothesis that DNA-directed termination downstream of protein-coding genes occurs when PAS cleavage cannot, Han *et al*., observed the most DNA-directed termination after depleting the PAS endonuclease, Ysh1 (Han et al. 2023).

In contrast to promoter-proximal regions, T-tracts are rarely used for termination within gene bodies. In budding yeast, SPT5 inhibits DNA-directed termination in purified systems and may function similarly in mammals (Han et al. 2023). In mammals, there is a “pausing zone” within 3kb of the TSS where premature transcriptional termination is frequent(Fong et al. 2022). Escape from this pausing zone requires SPT5 phosphorylation and the reversal of this state re-establishes termination competence beyond the PAS (Cortazar et al. 2019). Thus, SPT5 phosphorylation is high where DNA-directed termination is poor and low where it is frequent. We speculate that pausing zones associated with unphosphorylated SPT5 are also efficient termination zones where a DNA-dependent process is common. Other mechanisms that improve elongation, such as telescripting by U1 snRNA (Kaida et al. 2010; Mimoso and Adelman 2023), may also suppress this type of termination within long introns.

Our early work on transcriptional termination used the human β-globin gene as a model and studied its dedicated CoTC termination element. RNA transcribed from the CoTC sequence was proposed to be co-transcriptionally cleaved and enabled our original discovery of the role of XRN2 in RNAPII termination (West et al. 2004). We hypothesised that cleavage of CoTC RNA provided an entry site for XRN2 to access RNAPII-associated RNA, yet a responsible endonuclease was never discovered. We now suggest that termination occurs directly over the CoTC element, which is compatible with our previous findings that CoTC “activity” generates exosome substrates (Dye and Proudfoot 2001; West et al. 2006). Downstream of protein-coding genes, DNA-directed/CoTC termination may help evict RNAPII that is ill-configured for normal 3’ end processing or facilitate termination downstream of weak PAS’s that are not processed.

We speculate that short human protein-coding genes have more DNA-directed termination downstream of the PAS because RNAPII elongation complexes do not have sufficient distance to mature and acquire resistance to T-rich elements. In this respect, these genes resemble promoter-proximal transcripts. This fits the idea that elongation competence is established over an initial ∼3kb window, within which we show DNA-directed termination is common (Figure 4E). The much shorter length of budding yeast genes (∼1.5kb on average) might explain why DNA-directed termination is more general downstream of the PAS compared to in humans (Han et al. 2023). Finally, several *in vitro* studies have claimed that RNAPII transcriptional termination is independent of PAS cleavage (Orozco et al. 2002; Zhang et al. 2015). The efficiency of PAS cleavage is typically poor *in vitro* and transcription assays are usually performed on short DNA templates. Therefore, the previous observations of PAS-independent termination in such systems may reflect DNA-directed termination of RNAPII that cannot engage the torpedo mechanism due to defective PAS processing.

To conclude, we describe a mode of mammalian RNAPII termination driven by T-rich DNA sequences. Together with the known function of T-tracts in terminating prokaryotic RNAP and eukaryotic RNAPIII, our findings imply a conserved susceptibility to such termination. The maturation of RNAPII elongation complexes provides resistance to this mechanism. Given the need to transcribe for long distances in mammals, this may be why factor-dependent mechanisms have evolved to couple transcriptional termination of RNAPII to RNA processing.

## MATERIALS AND METHODS

### Cell lines and culture conditions

HCT116 and HeLa cells were cultured in Dulbeccos Modified Eagle Medium (DMEM) media supplemented with fetal calf serum (10%) and penicillin/streptomycin at 37°C with CO_2_ (5%). Transfections were performed using Jetprime (Polyplus). *INTS1-AID* and *dTAG-EXOSC3* cells were generated by CRISPR/Cas9-mediated non-homologous end joining using 1µg of an INTS1/EXOSC3-specific guide RNA (gRNA) plasmid and 1µg of repair PCR product transfected into sub-confluent 6-well plates. Media was refreshed 16hr later and cells were expanded into 100mm dishes 48hr later. These contained selective media including 10ug/ml Blasticidin (for dTAG-EXOSC3) or 800 ug/ml Neomycin/30 ug/ml Hygromycin (for INTS1-AID). After 10-14 days, single colonies were selected for screening by PCR for confirmation of genomic insertion. For *INTS1-AID* cells, TIR1 was subsequently integrated into the AAVS1 safe-harbour locus (addgene #72835) using a similar protocol, selecting for Puromycin at 1mg/ml. *XRN2-dTAG* (Estell et al. 2023) and *INTS11-dTAG* (Eaton et al. 2023) cells are described previously. Additional molecules were used at the following concentrations: auxin (500 mM), dTAGv-1 (1 µM), 4_S_U (500 µM).

### Western blotting analyses

1µg of GFP reporter plasmid was transfected into 6-well plates (∼50% confluent) of unmodified HCT116 parent cells for 16hr. Cells were then lysed in RIPA buffer (150mM NaCl, 1% NP40, 0.5% sodium deoxycholate, 0.1% SDS, 50mM Tris-HCl at pH 8, 5mM EDTA at pH 8) for 20mins on ice. Extracts were clarified by centrifugation (10 mins, 13000rpm) retaining the total protein supernatant.

### Antibodies

anti-GFP (RRID:AB_2749857), anti-Nucleolin (RRID:AB_300442), anti-CPSF73 (RRID:AB_1268249), anti-EXOSC3 (RRID:AB_2278183), anti-XRN2 (RRID:AB_873178), anti-INTS1 (RRID:AB_2127258), anti-HA (RRID:AB_390918).

### RNA isolation

Total RNA was extracted using 1ml Trizol according to the manufacturers’ protocol. RNA was treated with Turbo DNase for 1hr at 37°C. Nuclear RNA was isolated by pelleting cells in PBS, resuspending cells in 5ml hypotonic lysis buffer (10 mM Tris pH5.5, 10 mM NaCl, 2.5 mM MgCl_2_, 0.5% NP40), and then underlaying with 1ml of a 10% sucrose cushion. RNA was isolated from this pellet using Trizol.

### qRT-PCR analyses

1µg of RNA was reverse transcribed into cDNA using random hexamers and Protoscript II Reverse Transcriptase (NEB). cDNA was diluted to 50µl. qRT-PCR was performed using LUNA SYBR (NEB) on a Rotorgene (Qiagen). Fold changes were calculated using the ΔCT procedure. Further graphical analysis was achieved in R.

### 4sU labelling and biotinylation

HeLa cell data is published (Lykke-Andersen et al. 2021). For HCT116 cells, 2x 100mm culture dishes of cells were labelled with 4_S_U for 5mins. Total RNA was extracted, and DNase was treated as described above. For biotin labelling, 5µg of 4sU labelled Drosophila S2 4_S_U labelled RNA was spiked into 300µg extracted RNA samples and mixed with 10mM HEPES [pH 7.5], 1mM EDTA, and 10µg MTSEA biotin-XX (Biotium), 1µl RNase inhibitor to a final volume of 250µl. 50µl of dimethylformamide was added and samples were incubated at room temperature for 45mins in the dark. Samples were made up to 400µl with water, phenol-chloroform extracted, and ethanol precipitated. Pellets were resuspended in 200µl wash buffer (10mM Tris-Cl pH 7.4, 50mM NaCl, 1mM EDTA). 100µl of µMACS Streptavidin Microbeads (Miltenyi Biotec) were washed as previously described (Lugowski et al. 2018). 1x with nucleic acid equilibration buffer (provided) and 3x in 100µl wash buffer using a µMACS magnetic separator, before eluting the beads in 200µl wash buffer off the magnet and mixing with the resuspended RNA and 1µl RNase inhibitor. Columns were returned to the µMACS separator and washed 3x with 400µl wash 1 buffer (10mM Tris-Cl pH 7.4, 6M urea, 10mM EDTA) prewarmed to 65°C, and 3x with 400µL wash 2 buffer (10mM Tris-Cl pH 7.4, 1M NaCl, 10mM EDTA). RNA was eluted by 4x washes with 100mM dithiothreitol (DTT) dissolved in wash buffer and ethanol precipitated after incubating at −20°C for 1hr. Trace DTT was removed by precipitating again in 80% ethanol, and centrifuge at 13000rpm for 10mins.

### 3’-seq

*E. Coli* polyA polymerase (EPAP) treatment was performed using the Poly(A) Tailing Kit (Thermo) as previously described with some modification (Wu et al. 2020). 300ng of 4_S_U biotinylated RNA samples were divided into two and prepared for EPAP+ and EPAP-reactions containing 8µl 5x EPAP buffer, 4µl 25 mM MnCl_2_, 0.4µl EPAP, 0.4µl RNase inhibitor to a final volume of 40µl with water, replacing EPAP enzyme with water for -EPAP reactions. Samples were incubated for 30mins at 37°C, phenol-chloroform was extracted, and ethanol precipitated by centrifugation at 13000rpm for 10 mins. Next, rRNA was depleted using the NEBNext® rRNA Depletion Kit v2 (Human/Mouse/Rat) kit and libraries were generated using QuantSeq 3‘mRNA-Seq Library Prep Kit (Lexogen). Libraries were sequenced on an Illumina NovaSeq6000 (Novogene UK).

### POINT- and POINT5-seq

POINT-seq was performed as previously described (Sousa-Luis et al. 2021). Briefly, confluent 150mm plates were washed and scraped in PBS, pelleted before extracting nuclear RNA as above. Chromatin-associated RNAPII was extracted as follows. Nuclei were resuspended in 100µl NUN1 (20 mM Tris-HCl at pH 7.9, 75 mM NaCl, 0.5 mM EDTA, 50% glycerol, 0.85 mM DTT), before being incubated with 1 mL NUN2 +Empigen (20 mM HEPES at pH 7.6, 1 mM DTT, 7.5 mM MgCl_2_, 0.2 mM EDTA. 0.3 M NaCl, 1 M urea, 1% NP40, 3% Empigen) on ice for 15 min. The subsequent chromatin gel was washed in 10 ml PBS, resuspended in 250 µl DNase buffer (10mM Tris-HCl (pH 7.5), 400mM NaCl, 100 mM MnCl_2_,4 ml RNase inhibitor, and 12 µl Turbo DNase), and incubated at 37°C for 15 mins before centrifugation (10 mins, 13000rpm). The resulting supernatant was diluted 10-fold in NET2E buffer (50mM Tris-HCl (pH 7.4), 150mM NaCl, 0.05 % NP-40, and 3% Empigen) and precipitated at 4°C for 1.5hr with rotation using 10µg anti-RNAPII conjugated to 200µl sheep anti-mouse IgG Dynabeads (Thermo). The beads were washed 6x with ice-cold NET2E, before isolating RNA with Trizol. RNA was divided into two before processing into POINT and POINT5 libraries as described. POINT-seq libraries were made with the NEBNext® Ultra^TM^ II Directional RNA Library Prep Kit (NEB). POINT5-seq libraries were made with the SMARTer Stranded RNA-Seq kit (Takara Bio) protocol. Libraries were sequenced on an Illumina NovaSeq6000 (Novogene UK).

### Nuclear RNA-seq

Nuclear RNA-seq libraries were prepared from DNA-free nuclear RNA using NEBNext® Ultra^TM^ II Directional RNA Library Prep Kit (NEB) after first depleting rRNA using the NEBNext® rRNA Depletion Kit v2 (Human/Mouse/Rat). Sequencing was by Illumina NovaSeq6000 (Novogene UK).

### Primer sequences

Table S1.

### Illumina data pre-processing

Adapters were removed from raw Illumina reads using Trim_galore (https://github.com/FelixKrueger/TrimGalore(Martin 2011)) and aligned to Ensemble GRCh38 using Hisat2 (Kim et al. 2015) (default settings). Aligned reads were next filtered using SAMTools discarding unmapped, poorly mapped (MAPQ < 30), multi-mapped, and if paired-end, discordant reads (Li et al. 2009). Biological replicates were merged using BamTools after PCA plot verification (Barnett et al. 2011). Afterward, merged replicate samples were split by strand using SAMTools, and converted into RPKM normalised single nucleotide resolution bigwigs using DeepTools, scaling the antisense strand coverage by −1 (Ramirez et al. 2014). For +/−EPAP 3’-seq data, the 3’ end of the original aligned read was extracted using BedTools (Quinlan and Hall 2010). Bigwigs were additionally scaled using aligned S2 spike-in as a normalisation factor and normalised by RPKM. *INTS1-AID* POINT and POINT5 coverage was normalised to BPM (Bins Per Million). Additionally, for POINT5, the 5’ end of the aligned read was extracted, preserving its original directionality before the generation of normalised coverage. log_2_ change read coverage files were generated using the bigwigCompare function of DeepTools from single nucleotide strand-specific normalised bigwigs. Bigwigs were visualised using IGV (Robinson et al. 2011).

### Metagene analysis

Metagenes were generated using DeepTools computeMatrix from strand-specific normalised read coverage over a curated list of BED intervals with further graphical processing within R. In some cases, line smoothing was applied using the following formula: y ∼ s (x, bs = “cs”).

### snRNA 3’ box motif discovery

Starting with a list of HCT116 expressed snRNA genes, derived from untreated nuclear RNA-seq data, we performed MEME motif discovery using the initial 30 nucleotides immediately downstream of annotated snRNA TES. Genome-wide consensus motif scanning was performed using FIMO, extracting BED intervals intersecting core chromosomes only. Further analysis of FIMO box data was performed within R using the ChIPseeker package(Yu et al. 2015).

### PolyN termination analysis of +EPAP 3’-seq

PolyT/G/C hexamer sequences were extracted in bed format using the SeqKit locate function and a genomic fasta sequence file. Overlapping polyN hexamer intervals of the same orientation were merged before sub-dividing them into groups based on their overlap to in-house curated snRNAs, multi-exon TU, and mono-exonic PROMPTs interval lists using BedTools intersect. Heatmaps were produced using strand-specific log_2_ change (-EXOSC3/+EXOSC3) RPKM normalised 3’ end read coverage to each of the interval’s list, scaling the interval length to 10 nucleotides and including an upstream and downstream region of identical length.

### Identification of genes with exosome hypersensitive 3’ flanks

Starting with primary transcript isoforms of protein-coding genes, +/−EXOSC3 nuclear RNA-seq data was used to identify potential CoTC-type terminating transcripts by searching for those that fit the following criteria. The log_2_ change of coverage over the gene body remained ≤ 1, and the downstream 3 kb flanking region signal increased by ≥ 1.5 in the -EXOSC3 condition. A final list was then manually curated before further analysis.

### Genome Browser views

These were generated using the seqNdisplayR package (Lykke-Andersen et al. 2024).

### Definition of TU downstream regions

The downstream area of TUs was defined based on the HeLa annotation (Lykke-Andersen et al. 2021), using custom R scripts. For each strand, the distance between the end of a TU and the gene body of a downstream one was measured. TUs for which this distance was shorter than 100bp or overlapped with the end of chromosomes were filtered out. Over the defined distance, the rtracklayer package was used to measure sequencing coverage for each base of two replicates of RRP40 siRNA depletion TT-seq. A pseudo-count of 1 was then added to this coverage which was then log_2_-transformed and averaged between replicates. For each interval previously defined, a mean log coverage over the first 50bp downstream of the TES was measured, and intervals where it was below 2.5 were filtered out. To ensure that the downstream area would not encompass TUs invading downstream ones, an additional filtering was performed requiring coverage to reach 0 for at least 100 consecutive base pairs in the defined intervals. Finally, for each interval, the log coverage was binned by 50bp, and sudden drops of signal between bins were measured. The first position where the signal dropped by more than 2-fold between two consecutive bins was considered the end area. TUs for which such a drop could not be identified were filtered out. Following this procedure, a downstream area was defined for 10569 TUs out of the 23145 originally present in the annotation. Importantly, this subset of the annotation did not display any bias toward specific biotypes.

### Definition of exosome/XRN2-sensitive 3’ ends

These were defined using a custom R script. Using the previously defined downstream area, sequencing coverage for each base was measured using the rtracklayer package in relevant 3’-seq datasets treated with EPAP. A pseudo-count of 1 was applied at each position and log_2_-transformation was performed. For each replicate, we measured the log_2_(cov) difference between EXOSC/XRN2 depletion and controls. These Δlog_2_(cov) were then averaged. Stabilised 3’ends were defined as those displaying an average Δlog_2_(cov) above 0.5.

### PAS strength prediction

PAS strength prediction was performed using custom R and Python script and the APARENT package (https://github.com/johli/aparent), as in (Wu et al. 2020). For each analyzed TU, the sequence 70bp upstream of the TES was retrieved and searched for the presence of one or more PAS hexamers, according to the ranked list of top 100 hexamers used in humans from (Gruber et al. 2016). When more than one hexamer was identified, the best-ranking one was chosen, and when more than one hexamer with the best rank was identified, the one closest to the position 20bp upstream of the TES was chosen. An area of 205bp placing the beginning of the identified hexamer at position 71 was then defined. For TUs where no hexamer could be identified, the 205bp area was defined placing the TES at position 71. The obtained sequences were then analyzed with the APARENT package, using the ‘aparent_all_libs_resnet_no_clinvar_wt_ep_5_var_batch_size_inference_mode_no_drop.h5’ model, to predict the cleavage probability by the cleavage and polyadenylation machinery. APARENT provides two scores ranging from 0 to 1, predicting cleavage efficiency based on the PAS hexamer located in position 71 (‘narrow score’) or anywhere within the 205bp sequence provided (‘wide score’). Based on the score’s distribution over the HeLa annotation, and the fact that some TUs such as PROMPTs were not expected to get a proper PAS hexamer, TUs with a ‘Strong’ PAS were defined as those having a ‘wide score’ above or equal to 0.5 and TUs with a ‘Weak’ PAS as having a ‘wide score’ bellow 0.1.

### GEO accession numbers

New data was generated as part of this publication: all HCT116 cell 3’-seq (GSE264499), HCT116 *INTS1-AID* cells POINT5-seq (GSE264716), HCT116 *INTS1-AID* cells POINT-seq (GSE264717) and HCT116 *dTAG-EXOSC3* cells nuclear RNA-seq (GSE264718). Published data was downloaded for: *DIS3* PAR-CLIP (GSE64332)*, XRN2-AID* POINT- and POINT5-seq (GSE159326), INTS11, RRP40, and INTS11xRRP40 siRNA depletion 3’-seq EPAP and non-EPAP treated (GSE151919), INTS11 and RRP40 siRNA depletion TT-seq and RNA-seq (GSE151919), XRN2-mAID nuclear RNA-seq and mNET-seq (GSE109003) and CPSF30-mAID nuclear RNA-seq (GSE163015).

## ACKNOWLEDGEMENTS

We are grateful to the Wellcome Trust for supporting this work with an Investigator Award 223106/Z/21/Z to S.W. Research in the N.J.P lab was also funded by a Wellcome Trust Investigator Award (107928/Z/15/Z). Research in the T.H.J. lab was supported by the Danish Cancer Society, the Lundbeck Foundation, and the Novo Nordisk Foundation (NNF) (ExoAdapt grant 31199). The project used the University of Exeter Sequencing Service funded by a Wellcome Trust multi-user equipment grant (218247/Z/19/Z). We thank Hiroshi Kimura for the RNAPII antibody employed for POINT analyses. Figure 7 was made using Biorender.

## AUTHOR CONTRIBUTIONS

Conceptualization, data curation, formal analysis, L.D., J.O.R., R.L-S, T.H.J., and S.W; Methodology, L.D., J.O.R., T.N. Investigation, L.D., J.O.R., and S.W; supervision and funding acquisition, N.J.P., T.H.J and S.W.; writing and editing, all authors.

---

**FIGURE S1.**
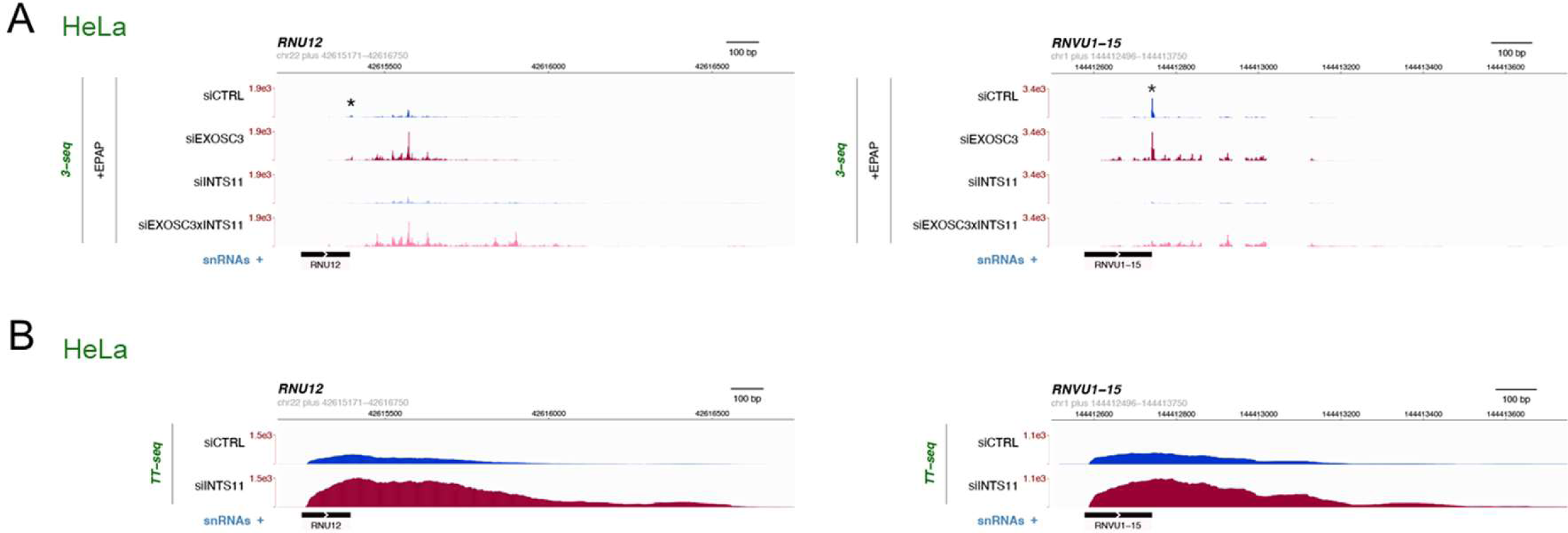
related to Figure 1. A. Genome browser views as in Figure 1B but displaying EPAP data across and downstream of the *RNU12* and *RNVU1-15* TUs. The data are from GSE151919 (Lykke-Andersen et al. 2021) B. Genome browser view as in (A) but displaying TT-seq data from siCTRL and siINTS11-treated HeLa cells (GSE151919 (Lykke-Andersen et al. 2021)).

**FIGURE S2.**
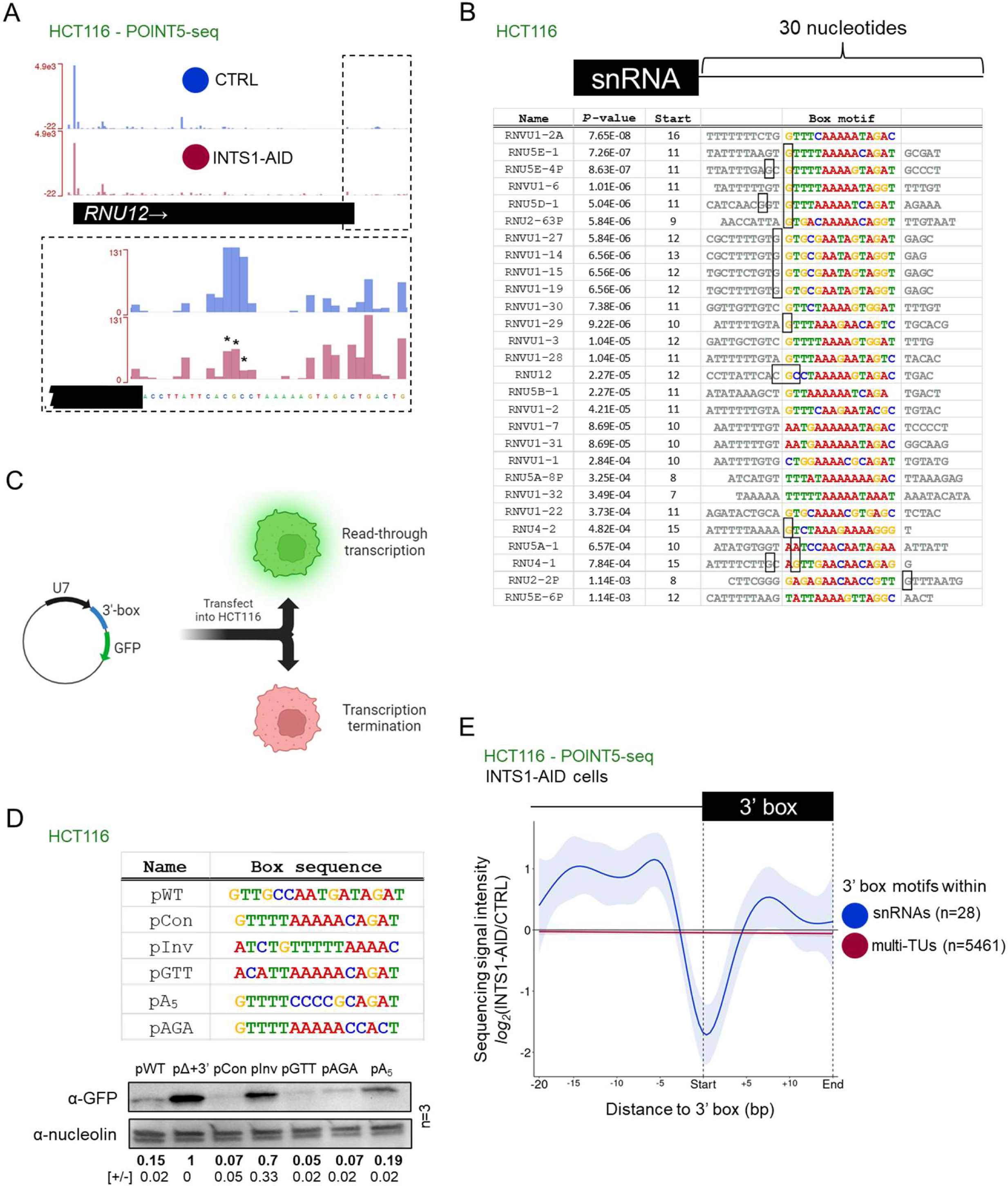
related to Figure 2. A. Genome browser view as in Figure 2D, but displaying POINT5-seq coverage over the *RNU12* TU. B. Result of MEME motif consensus over the first 30 nucleotides downstream of the TESs of the indicated Ensembl-annotated expressed snRNAs. The ‘start’ column indicates the number of nucleotides between the TES of each snRNA and the beginning of the 3’ box element (coloured). The first nucleotides downstream of significant POINT5-detectable and INTS1-sensitive cleavage sites are boxed. The p-value represents the confidence for each identified 3’ box motif. C. Schematic representation of the U7-GFP reporter plasmid (left) and assay (right). If processing occurs at the 3’ box, GFP expression is lost (red cell). Processing failure results in GFP expression (green cell). D. Western blotting analysis (bottom panel) of HCT116 cells transfected with GFP reporter constructs harbouring the indicated 3’ box motif variants (top panel). The mean relative GFP protein quantities were calculated relative to those obtained in pΔ3’ box samples (reporter variant lacking a 3’ box) after normalising to nucleolin protein levels (n=3, +/−values = SEM). E. Metaplot of POINT5-seq data from *INTS1-AID* HCT116 cells untreated (CTRL) or treated (INTS1-AID) with auxin. The x-axis covers 20 nucleotides upstream of the start of the 3’ box until the end of the 3’ box element. The y-axis shows the log_2_ signal intensity ratio (INTS1-AID/CTRL). The blue line shows data from 3’ box instances at snRNAs and the reduced signal at the start of the 3’ box signifies co-transcriptional processing by INT. The red line shows instances of the consensus 3’ box in the promoter-proximal regions of multi-exonic TUs and is unchanged by INTS1 depletion indicating no detectable INT cleavage activity.

**FIGURE S3.**
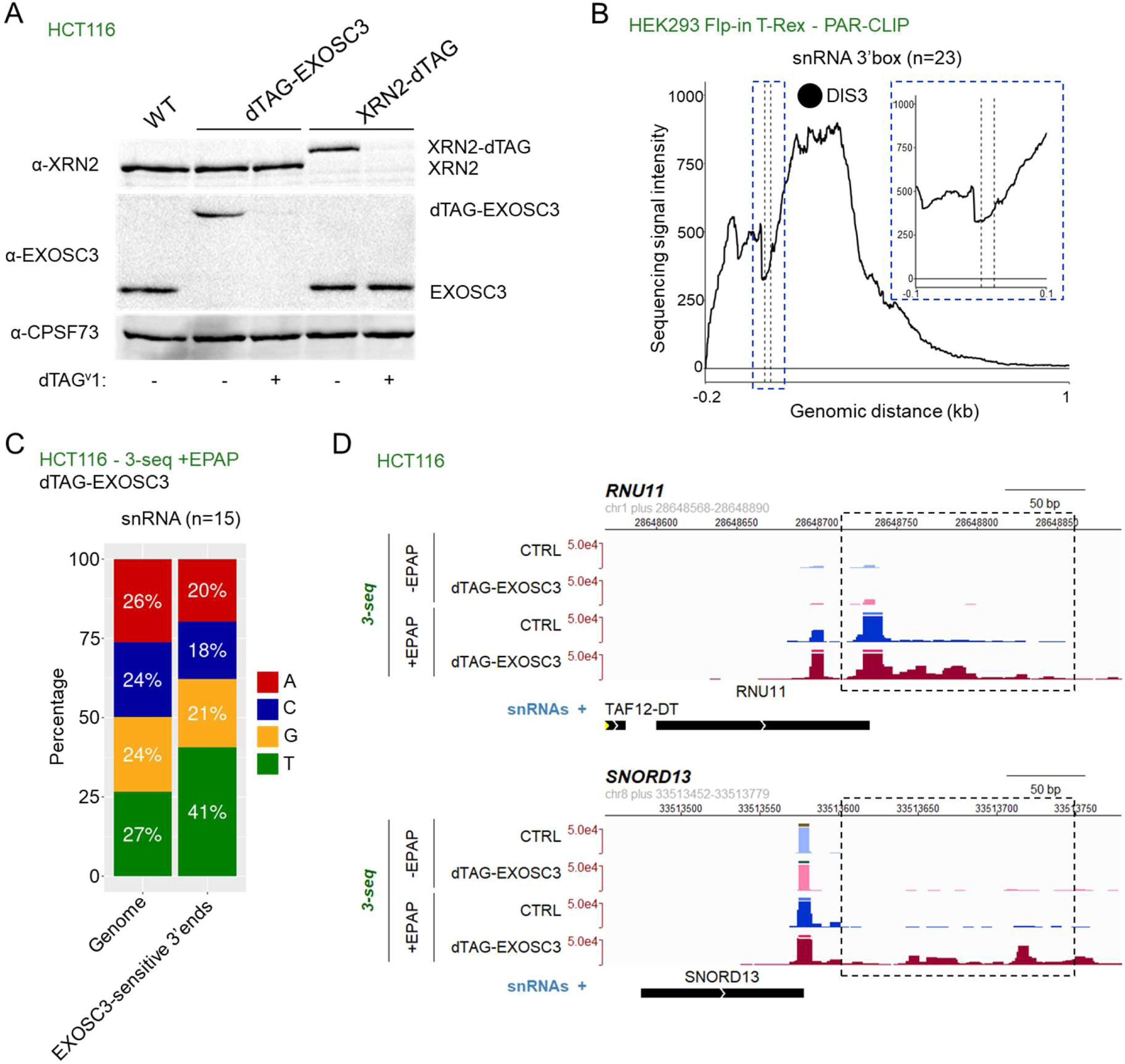
related to Figure 3. A. Western blotting analysis of HCT116, *dTAG-EXOSC3,* and *XRN2-dTAG*, cells treated, or not, for 4hr with dTAG-v1 as indicated beneath the blot. CPSF73 was probed as a loading control. Homozygous tagging of EXOSC3 and XRN2 was evident from the slower migration of the dTAG-EXOSC3 and XRN2-dTAG species, which were depleted following dTAGv-1 treatment. B. Metaplot of DIS3 PAR-CLIP data (GSE64332 (Szczepinska et al. 2015)) across the 3’ box region of 23 snRNAs. The x-axis covers the 3’ box (black dashed lines) as well as sequences encompassing 0.2kb upstream and 1kb downstream. The y-axis sequence signal intensity units are RPKM. The zoomed area (blue dashed box) highlights the 3’ box and the immediate increase in the downstream PAR-CLIP signal. C. Histogram showing the % of each nucleotide in the genomic sequence downstream of snRNA TESs (“Genome”) compared to the % of each nucleotide at the 3’ end of EPAP-treated 3’-seq reads over the same region that were stabilised by log_2_ change ≥1 in dTAGv-1 treated *dTAG-EXOSC3* HCT116 cells *vs.* CTRL HCT116 cells (“EXOSC3-sensitive 3’ ends”). D. Genome browser views of 3’-seq data obtained from -/+EPAP-treated RNA samples isolated from dTAG-treated dTAG-EXOSC3 cells or HCT116 CTRL cells. The *RNU11* and *SNORD13* TUs and 200 nucleotides of the downstream region are shown. The dashed boxes highlight clusters of 3’-seq reads that are EPAP-dependent and stabilised by dTAG-EXOSC3 depletion.

**FIGURE S4.**
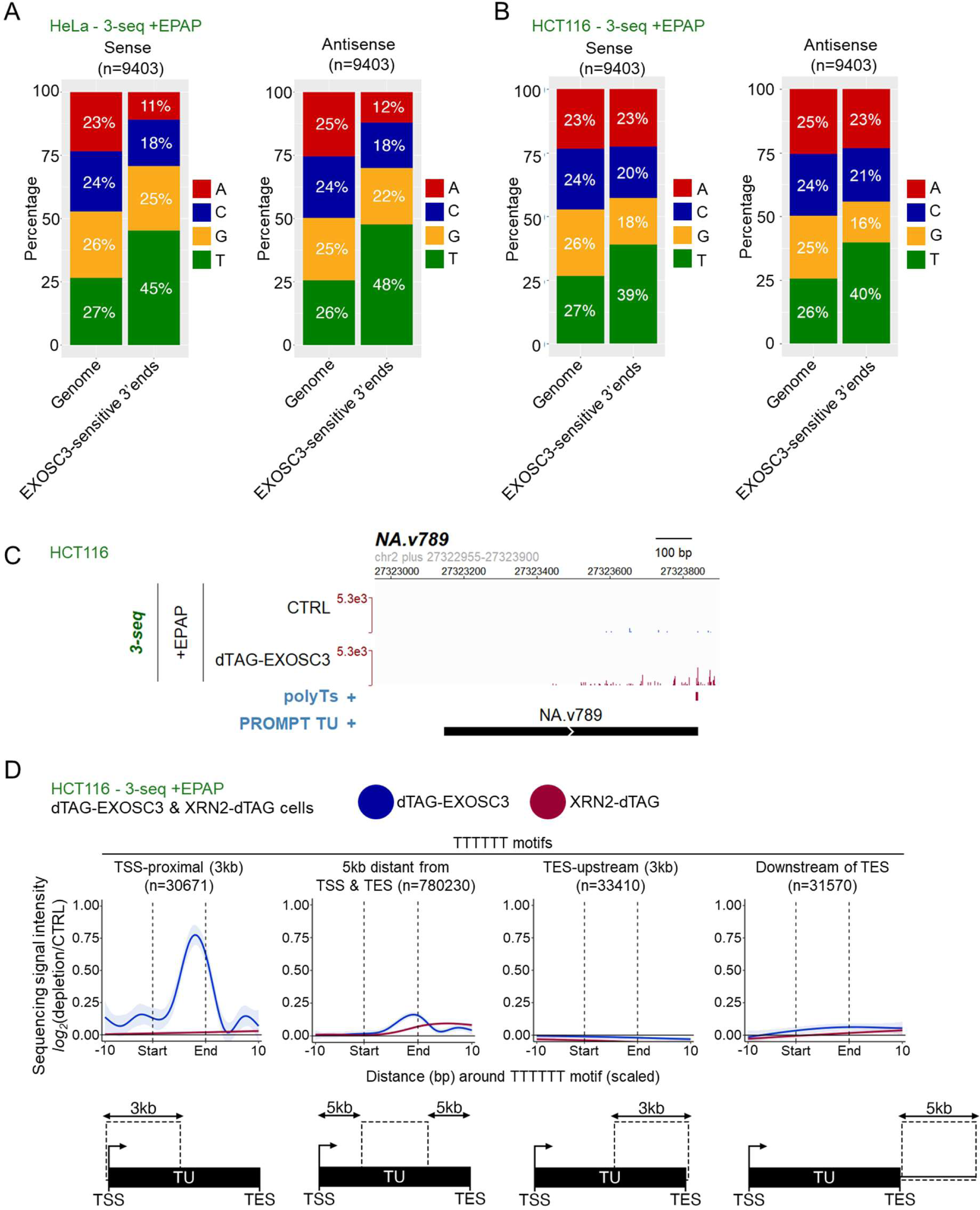
related to Figure 4. A. Histogram as in Figure S3C but showing the genomic sequence within 3kb sense (left) and antisense (right) of TSSs compared to the % of each nucleotide at the 3’ end of EPAP-treated 3’-seq reads over the same region that were stabilised by log_2_ change ≥1 in siEXOSC3-vs siCTRL treated HeLa cells (“EXOSC3-sensitive 3’ ends”). The data are from GSE151919 (Lykke-Andersen et al. 2021) B. Histogram as in (A) but for data obtained in dTAGv-1 treated *dTAG-EXOSC3* HCT116 cells *vs.* CTRL HCT116 cells. C. Genome browser view of a PROMPT TU showing +EPAP 3’-seq coverage in CTRL HCT116 cells and dTAG-EXOSC3 HCT116 cells treated with dTAGv-1 (4hr). The location of the T-tract is indicated by the red bar under the lower track. D. Metaplots showing the log_2_ sequencing intensity change in +EPAP 3’ end signals, comparing dTAG-treated *dTAG-EXOSC3* or *XRN2-dTAG* HCT116 cell samples with CTRL HCT116 cell samples. The plot displays multi-exonic TUs divided into four regions as indicated below: 3kb TSS-proximal, gene body (omitting the 5kb TSS- and TES-proximal regions, 3kb upstream of the TES, and 5kb downstream of the TES. Each plot displays signal changes across T≥6 sequences and includes 10 nucleotides upstream and downstream.

**FIGURE S5.**
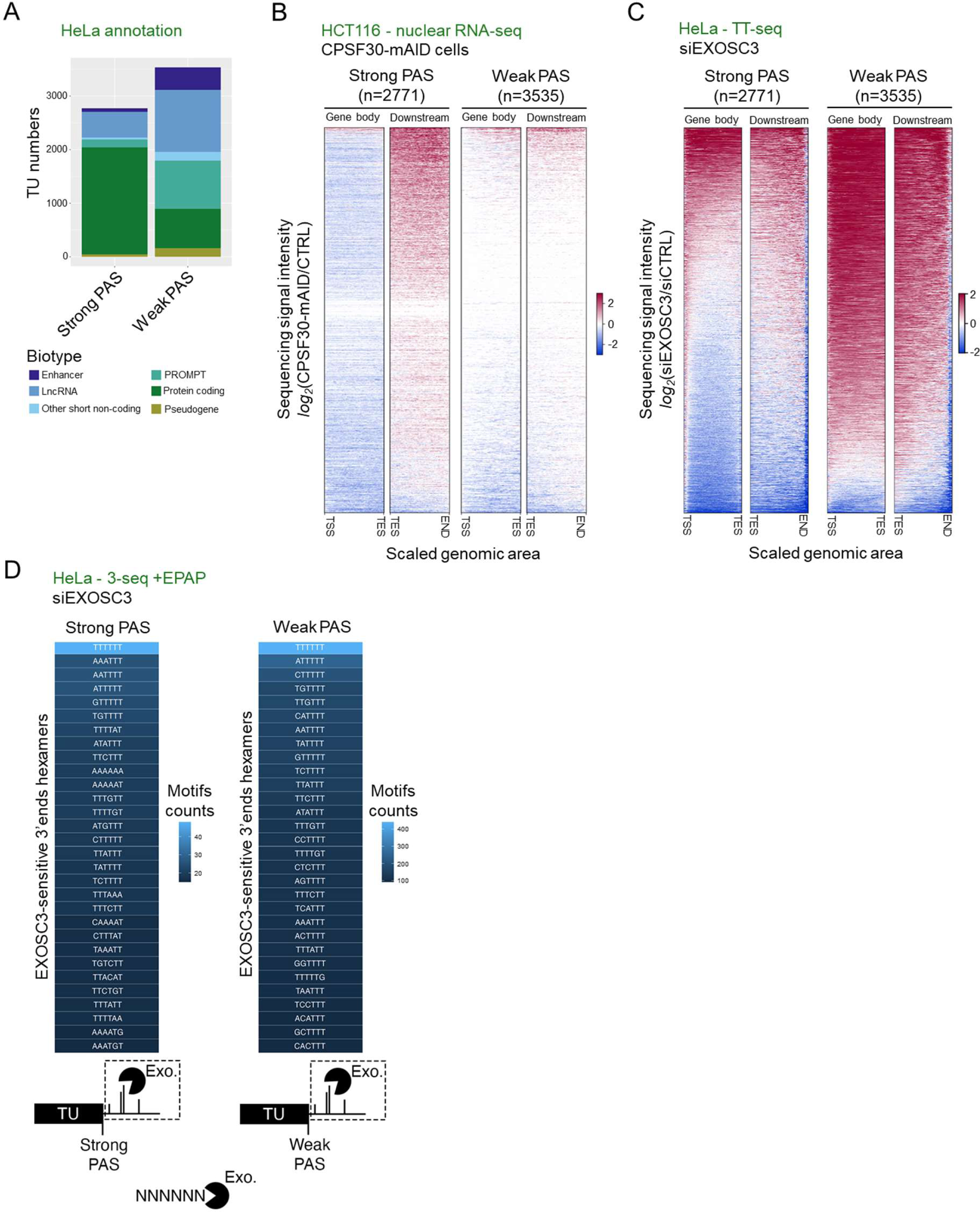
related to Figure 5. A. Histogram showing the proportion of the indicated RNA biotypes classified by APPARENT2 as having a strong or a weak PAS. All TUs with a 3’ end defined by TT-seq data after EXOSC3 depletion from HeLa cells were included (see Materials and Methods). B. Heatmap analysis as in Figure 5B but showing the log_2_ change in nuclear RNA-seq signal intensity following CPSF30-mAID depletion from HCT116 cells (CPSF30-mAID *vs.* CTRL from GSE163015 (Estell et al. 2021)). Note that CPSF30 loss increases signal downstream of strong but not weak PASs. CPSF30 loss also reduces the gene body signal upstream of strong PASs -a well-described consequence of rapid CPA factor loss (Eaton et al. 2020; Cugusi et al. 2022). C. Heatmap analysis as in (B) but for TT-seq data derived from siEXOSC3- and siCTRL-treated HeLa cells (GSE151919 (Lykke-Andersen et al. 2021)). D. Sequence composition analysis as in Figure 5D, but of 3’ ends beyond strong or weak PASs and stabilised by log_2_ change ≥1 following siEXOSC3 *vs*. siCTRL treatment of HeLa cells (GSE151919 (Lykke-Andersen et al. 2021)).

**FIGURE S6.**
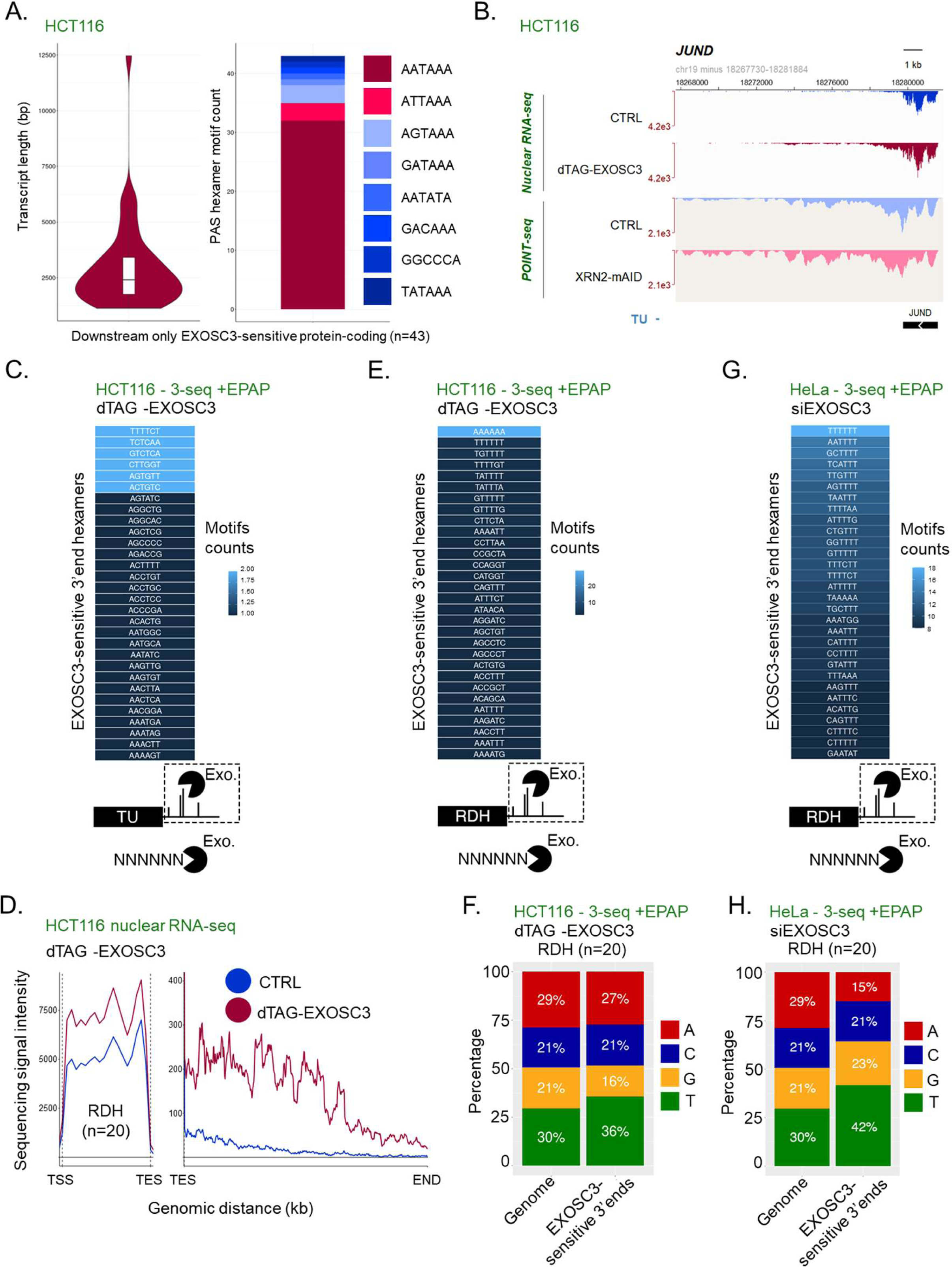
related to figure 6. A. Left: Violin plot showing TU length distributions for the 43 examples studied. The y-axis shows transcript length in nucleotides. Right: PAS hexamer motif counts for the 43 TUs. B. Genome browser view of *JUND* (one of the 43 TUs, harboring EXOSC3-sensitive 3’ flanking RNA). Nuclear RNA-seq-(top two tracks) or POINTseq-(bottom two tracks) signals from CTRL HCT116 cells or degron derivatives depleted of dTAG-EXOSC3 (nuclear RNA-seq) or XRN2-mAID (POINT-seq -GSE159326 (Sousa-Luis et al. 2021)) are shown. The y-axis scales are RPKM. C. Sequence composition analysis as in Figure 5D, but of 3’ ends deriving from downstream the TESs of the 43 studied TUs. D. Metaplots of nuclear RNA-seq signal from *dTAG-EXOSC3* HCT116 cells treated, or not, for 4hr with dTAGv-1. The left and right plots display signals over RDH gene bodies and downstream of RDH TESs, respectively. The y-axis sequence signal intensity units are RPKM. E. Sequence composition analysis as in (C), but of 3’ ends deriving from downstream RDH TESs. Note that although A6 is the top motif, this should be interpreted cautiously due to the employed EPAP activity. Other highly enriched motifs are T-rich. F. Histogram as in Figure S3C but showing the % of each nucleotide in the genomic sequence downstream of RDH TESs (“Genome”) compared to the % of each nucleotide at the 3’ end of EPAP-treated 3’-seq reads over the same region that were stabilised by log_2_ change ≥1 in dTAGv-1 treated *dTAG-EXOSC3* HCT116 cells *vs.* CTRL HCT116 cells (“EXOSC3-sensitive 3’ ends”). G. Sequence composition analysis as in (E), but displaying signals derived from siEXOSC3 *vs.* siCTRL-treated HeLa cells (GSE151919 (Lykke-Andersen et al. 2021)). H. Histogram as in (F) but displaying samples derived from siEXOSC3 *vs.* siCTRL-treated HeLa cells.

